# Single-particle multi-parametric microscopy reveals structural, size, and payload heterogeneity in mRNA-loaded lipid nanoparticles

**DOI:** 10.1101/2025.10.17.683130

**Authors:** Albert Kamanzi, Ariadne Tuckmantel Bido, Yao Zhang, Erik Olsén, Maya Stibbards-Lyle, Martin Jasinski, Yifei Gu, Benjamin Wang, Michael Venier-Karzis, Romain Berti, Michelle Jeliazkova, Cynthia Shaheen, Jerry Leung, Miffy Hok Yan Cheng, Pieter Cullis, Sabrina Leslie

**Affiliations:** Michael Smith Laboratories, University of British Columbia, Vancouver, BC, V6T 1Z4, Canada; Department of Physics and Astronomy, University of British Columbia, Vancouver, BC, V6T 1Z1, Canada; ScopeSys, Inc., 2366 Main Mall, Vancouver, BC, V6T 1Z4, Canada; School of Biomedical Engineering, University of British Columbia, Vancouver, BC, V6T 2B9, Canada; Department of Biochemistry and Molecular Biology, University of British Columbia, Vancouver, BC, V6T 2A1, Canada; Department of Physics, University of Gothenburg, Gothenburg, 41296, Sweden; Faculty of Pharmaceutical Sciences, University of British Columbia, Vancouver, BC, V6T 1Z3, Canada

**Keywords:** lipid nanoparticles, mRNA payload, single-molecule microscopy, drug delivery, FRET, nanomedicines, structure function relationship, microfluidics, biophysics

## Abstract

Deciphering the heterogeneity of mRNA-containing lipid nanoparticles (LNPs) is essential for understanding the relationship between their microscopic properties and therapeutic function. Here, by combining alternating laser excitation (ALEX) with Convex Lens-induced Confinement (CLiC) microscopy, we simultaneously measure size, multi-color fluorescence, mRNA payload, and Förster resonance energy transfer (FRET) of individual suspended LNPs containing labeled lipid and mRNA molecules. By varying formulation parameters, including ionizable lipids, formulation buffers, and molecular ratios, we investigated and correlated key microscopic properties for relevant vaccine formulations. While the per-particle lipid fluorescence was lower for empty versus mRNA-loaded particles for all formulations, the relative size of empty versus mRNA-loaded particles depended upon the formulation and intraparticle structure. When comparing CLiC-ALEX to cryogenic transmission electron microscopy measurements (Cryo-TEM), for the LNP formulations that display blebs, the fraction of bleb LNPs is in close agreement with the fraction of mRNA-containing LNPs. CLiC-ALEX also enabled quantification of the per-particle mRNA fluorescence and FRET signals, and thus the heterogeneity in the mRNA copy number and mRNA-LNP structural arrangements, where the results were compared with biophysical estimates based on the LNP formulations. These rigorous biophysical insights are critical to inform our understanding of structure-activity relationships and inform rational design of nanomedicines.

---

Lipid nanoparticles (LNPs) are widely used to deliver mRNA-based therapies, ^1^ and their potential was highlighted by the rapid response to the COVID-19 pandemic with the Tozinameran (Comirnaty^®^)^2^ and Elasomeran (Spikevax^®^)^3^ vaccines. Despite this success, it remains a challenge to further optimize LNP performance, particularly to improve delivery efficiency and to enable targeting of tissues beyond the liver. ^4^

An aspect of this challenge comes from the fact that different LNP formulations and manufacturing procedures significantly affect the size and Morphology of the LNP, characterized by spherical and nonspherical shapes,^5,6^ which in turn affects their therapeutic function.^7^ For example, recent studies have shown that nonspherical LNPs containing ‘blebs’ exhibit enhanced transfection efficiency compared to spherical LNPs. ^5^ Other examples include liposomal particles formulated with an increased proportion of bilayer lipids,^8^ which demonstrated prolonged circulation lifetimes for extrahepatic delivery. As a result, the observed correlation between mRNA-LNP structure and their therapeutic performance has raised the need for detailed characterization of mRNA-LNPs during development and manufacturing.

In addition to the major variations in size and structure between different formulations, there are often multiple subpopulations of particles within the same formulations, such as LNPs with and without mRNA payloads or with and without blebs,^9^ which are not captured by commonly used ensemble-average measurements. Here, we refer to LNPs containing one or more mRNA molecules as loaded and those without mRNA as empty LNPs. Ensemble-average measurements, such as encapsulation efficiency and particle size, are commonly used by developers for LNP quality assessment^10^ but do not distinguish between loaded and empty LNPs. To be able to relate particle heterogeneity to function, measurements need to not only accurately capture particle size, loading, and structure information but also capture this on the single-particle level under physiologically relevant conditions.

Multiparametric single LNP characterization is often based on fluorescence, either fluorescence alone or combined with simultaneous light scattering measurements, where labeled cargo is used in combination with a fluorescent lipid.^9,12–16^ For example, the fraction of empty LNPs has previously been measured by imaging LNPs that contain fluorescent lipids and labeled mRNA molecules. ^9,12,13,15,–17^ Combined with simultaneous size measurements, the scaling between signal and size provides structural information.^15,18^

In addition to investigating the fraction of loaded LNPs, when negligible coupling between fluorophores is assumed, scaling the cargo fluorescence signal with the signal of a single cargo molecule can be used to estimate the number of mRNA molecules per particle.^12,13^ However, coupling between fluorophores within LNPs can, for example, cause Förster resonance energy transfer (FRET), where FRET has been both a source of signal uncertainty^13^ and a tool for probing LNP integrity.^19^ Thus, to reduce the uncertainty in estimating the amount of mRNA per LNP and provide information on the molecular arrangement within an LNP, simultaneous FRET quantification is required.

In this work, we address the need for a deeper understanding of LNP heterogeneity by using high-throughput single-particle measurements of size, payload, and molecular arrangements made under suspended-solution conditions. We achieve this by combining two-color Convex Lens-induced Confinement (CLiC)^20–22^ microscopy and alternating laser excitation (ALEX), enabling both sequential and simultaneous imaging of direct fluorescence emissions across two channels from individual particles, as well as their interactions through FRET. CLiC enables imaging of the extended time courses of many single diffusing particles that are isolated and confined to the focal plane of the microscope. ^18^ Using a series of clinically relevant mRNA-LNP formulations representing approved vaccines and other promising nanomedicines candidates (Table 1), we investigate the correlated distribution of size, as well as lipid, mRNA loading, and FRET signals, and compare our observations to cryogenic transmission electron microscopy (Cryo-TEM) and biophysical estimates based on the LNP formulations. This enables a detailed and direct comparison of similarities and differences across formulations to better understand the underlying heterogeneity of the nanomedicines.

**Table 1:**
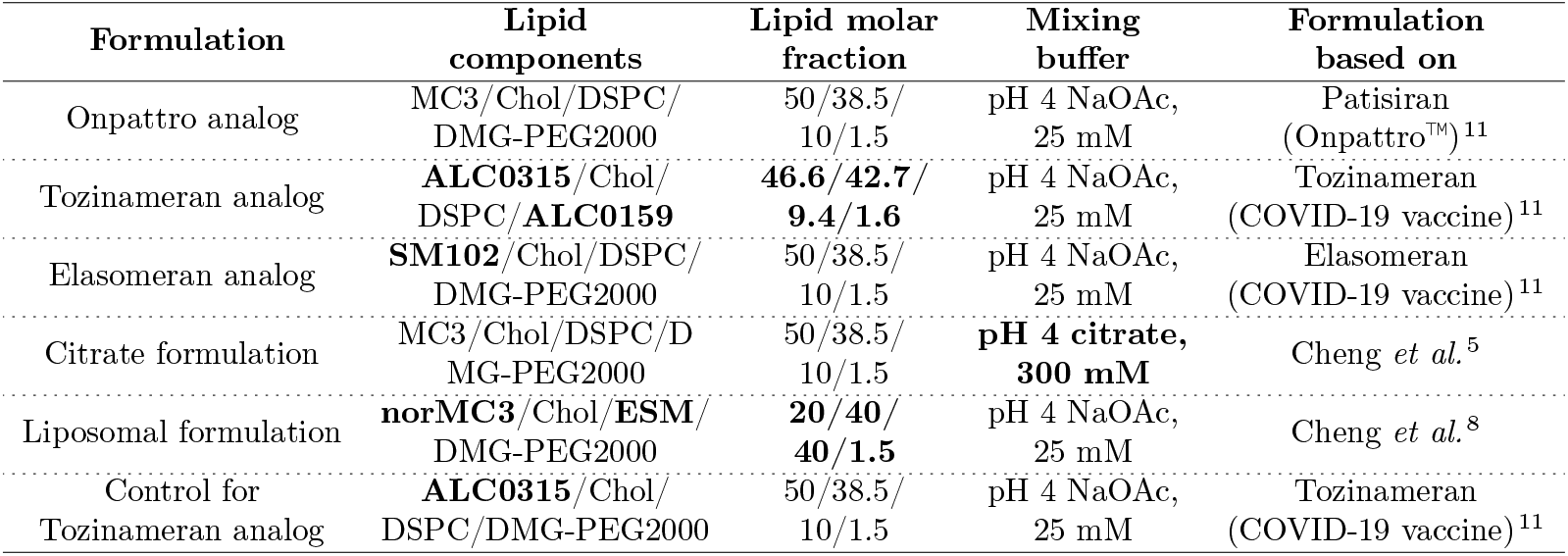
A list of formulations used, which were chosen to match/represent approved vaccines and medicines, or recent publications on LNP formulation design. All the formulations used were loaded with Cy5 Firefly Luciferase mRNA (1921 nucleotides), at an N/P ratio of 6. The top row shows a formulation based on Patisiran (Onpattro^®^) drugs, while the second and third row are analogs of Tozinameran (Comirnaty^®^) and Elasomeran (Spikevax^®^) COVID19 vaccines respectively. Both row 4 and 5 are based on recently published work, in which therapeutic activity increased in animal studies as a function of changes in formulation parameters. The last formulation was added in as a control for the Tozinameran (COMINARTY™) vaccine analog, as it uses a different PEG-lipid as well as lipid mixing fraction. The text in bold highlights differences among the formulations relative to the Onpattro analog, which included various ionizable lipids, lipid-mixing fractions, and formulation buffers.

## Results and Discussion

### CLiC-ALEX overview

To enable combined simultaneous and sequential microscopy of individual suspended LNPs for detailed size and fluorescence quantification, we combine CLiC single-particle microscopy with two excitation lasers (488 nm and 647 nm wavelengths) and a two-camera configuration, where we control the combination of lasers used for each frame with an acousto-optic tunable filter (AOTF) (Figure 1A). The CLiC device’s imaging flowcell uses coverslips containing an embedded array of microwells in which each microwell is 500 nm deep, matching the focal depth, and 3 µm in diameter, where all surfaces in the flow cell are passivated using PEGylation (see Methods, “Imaging Flow-cell Cleaning, Surface Treatment, and Assembly”).^18^ Using the CLiC device, we load LNP samples into a deflectable flow-cell chamber (Figure 1B(i)) and confine individual LNPs in microwell features embedded in the bottom surface (Figure 1B(ii, iii)) by deflecting the top surface into contact with the bottom. Particle tracking is then used to obtain trajectories of individual particles. From each trajectory, a mean squared displacement curve is generated (Figure 1B(iv)), which is fit to a particle-confinement model to determine particle diffusivity.^18,23,24^ The diffusivity is then converted to the hydrodynamic radius. ^18,25^

**Figure 1:**
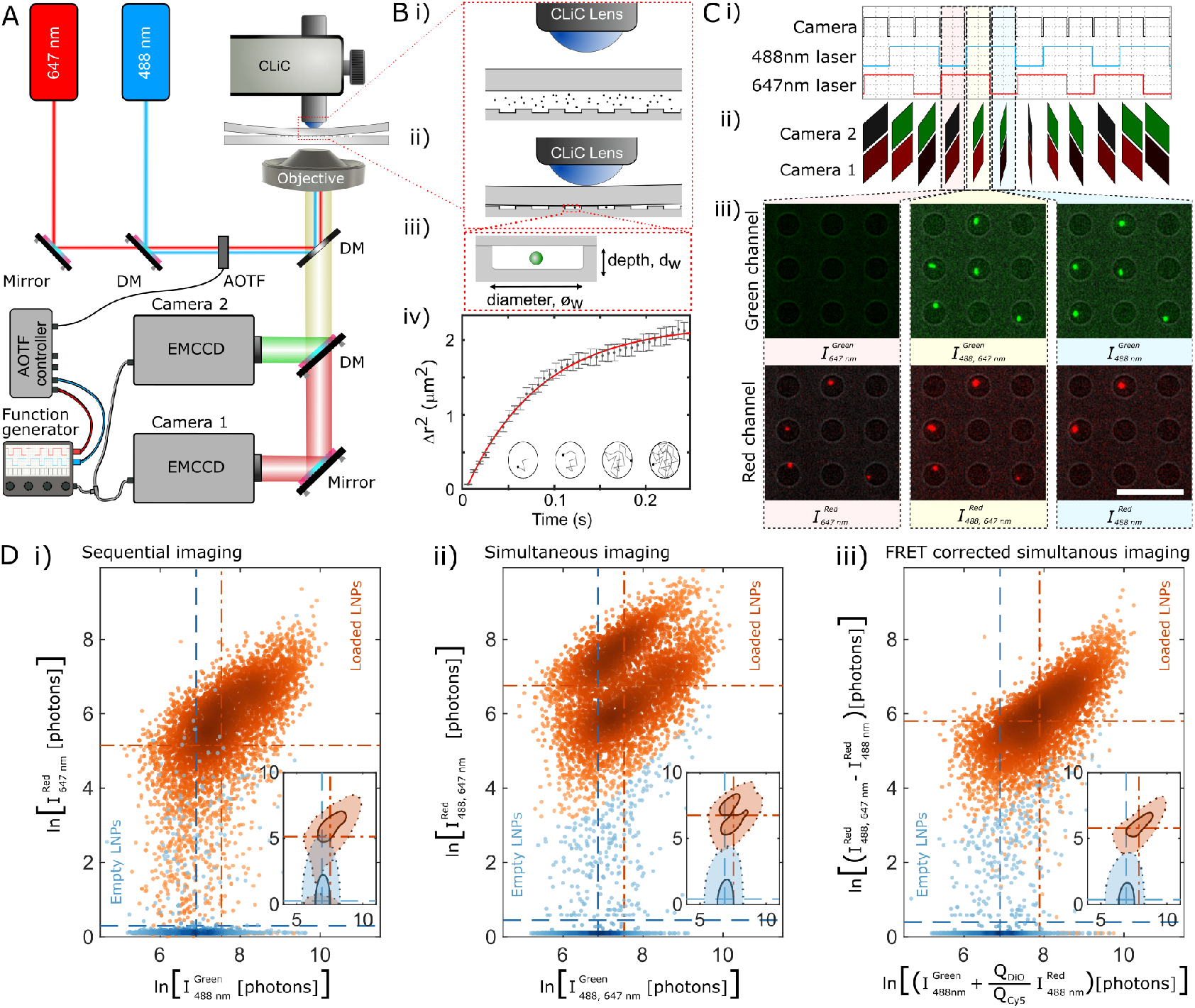
Single-particle CLiC-ALEX microscopy allow for improved quantification of signals from multiple fluorophore species on individual particles, and their interactions: (A) Schematic of the imaging system, including: a two-channel alternating laser excitation setup, a CLiC device on an inverted fluorescence microscope, and a dual-camera system for data collection. Laser timing is controlled by an Acousto-optic tunable filter (AOTF) and synchronized with the camera frame times by using a function generator as the trigger source. The CLiC device includes a deflectable flow-cell chamber and a pusher lens, as shown in (B i., ii.). This allows for trapping of individual freely diffusing LNPs in CLiC micro-wells (B iii.), which are then tracked to generate long trajectories for mean squared displacement (MSD) calculations (B iv.). The error bars in the MSD plot represent the standard deviation values in the MSD values at each time point. (C i.) Triggering waveforms are generated by the function generator, to control both cameras (black line), as well as the 488-nm (blue line) and the 647-nm (red line) lasers. By triggering the lasers at a frequency three times lower than the camera frame rate, with a phase delay of one exposure time, we are able to combine both sequential and simultaneous excitations across our imaging channels, as shown in C (ii., iii.). (C iii.) Example images of LNPs as observed in the two channels under the three excitation conditions, Scale bar = 10 µm. Here, fluorescent images are false colored, to represent imaging channels, and are overlayed with a bright field image to indicate location of CLiC micro-wells. However, no additional processing (*e.g*., background subtraction) was performed. (D) The resulting signal quantification using different excitation modalities, including (D i.) sequential excitation, with either the red or blue laser on (pink or blue band in C, respectively), (D ii.) simultaneous excitation, with both lasers on (yellow band in C) and (D iii.) a specific superposition of intensities (to segment one signal cluster). Here we note that it is challenging to distinguish between empty and loaded LNPs using sequential excitation, as tracking of the Cy5 signal is noise sensitive. This is resolved by simultaneous excitation due to position-assisted averaging (see Methods, “Colocalization classification”), however, simultaneous excitation is instead biased by dye-dye interactions. By combining both methods we are able to distinguishing between loaded and empty LNPs while also accurately quantifying the LNP-DiO, mRNA-Cy5 and FRET signals on the single-particle level.

Using the CLiC-ALEX method, we investigate LNPs loaded with mRNA molecules as a function of key formulation parameters (Table 1), in which the mRNA molecules are covalently labeled with Cy5 fluorophores (647/670 nm) and their lipid carriers include DiO (DiOC18(3) (3,3’-Dioctadecyloxacarbocyanine Perchlorate), 488/505 nm) lipophilic dyes. For this work, we selected formulations containing ionizable cationic lipids (one of MC3, ALC0315, SM102 or norMC3), cholesterol (Chol), distearoylphosphatidylcholine (DSPC), a PEG-lipid (DMG-PEG2000 or ALC0159), and lipophilic dyes (DiO), mixed in varying both lipid molar ratios and preparation buffers (Methods, “Preparation of the lipid nanoparticles” and Table 1).

To enable ALEX, we trigger the alternating lasers synchronously with the cameras (Figure 1C), while allowing for both sequential and simultaneous single-particle signal measurements (Figure 1D (i, ii)). This allows direct estimation of single-particle fluorescence and FRET by comparing the difference in the measured fluorescence intensity during simultaneous imaging and sequential imaging (Methods, “Fluorescence signal quantification”). By correcting for the FRET contribution, as shown in Figure 1D (iii), the secondary cluster from Figure 1D (ii) disappeared. Combined with particle tracking size estimation (Methods, “CLiC particle sizing”) and fluorescence signal colocalization analysis (Methods, “Colocalization classification”), this allows us to accurately quantify direct signals from DiO and Cy5, as well as their interaction through FRET at the single-particle level, where simultaneous imaging enables a clear signal distinction between loaded and empty LNPs.

### Formulation-dependent differences in the clustering of loaded and empty LNPs

When investigating the relationship between DiO intensities (*I*_DiO_) and the corresponding hydrodynamic radii (*r*_h_) for the six formulations listed in Table 1, we observe that loaded LNPs (mRNA-positive) are generally brighter but not necessarily larger than empty ones (mRNA-negative), as shown in Figure 2. Loaded LNPs have an average radii of 32-44 nm versus 26-34 nm for empty particles, while their average DiO intensities range from 2850-7640 photons compared to 1310-1700 photons. Thus, variation in the DiO signal across formulations and loading conditions is greater than the corresponding variation in particle size.

**Figure 2:**
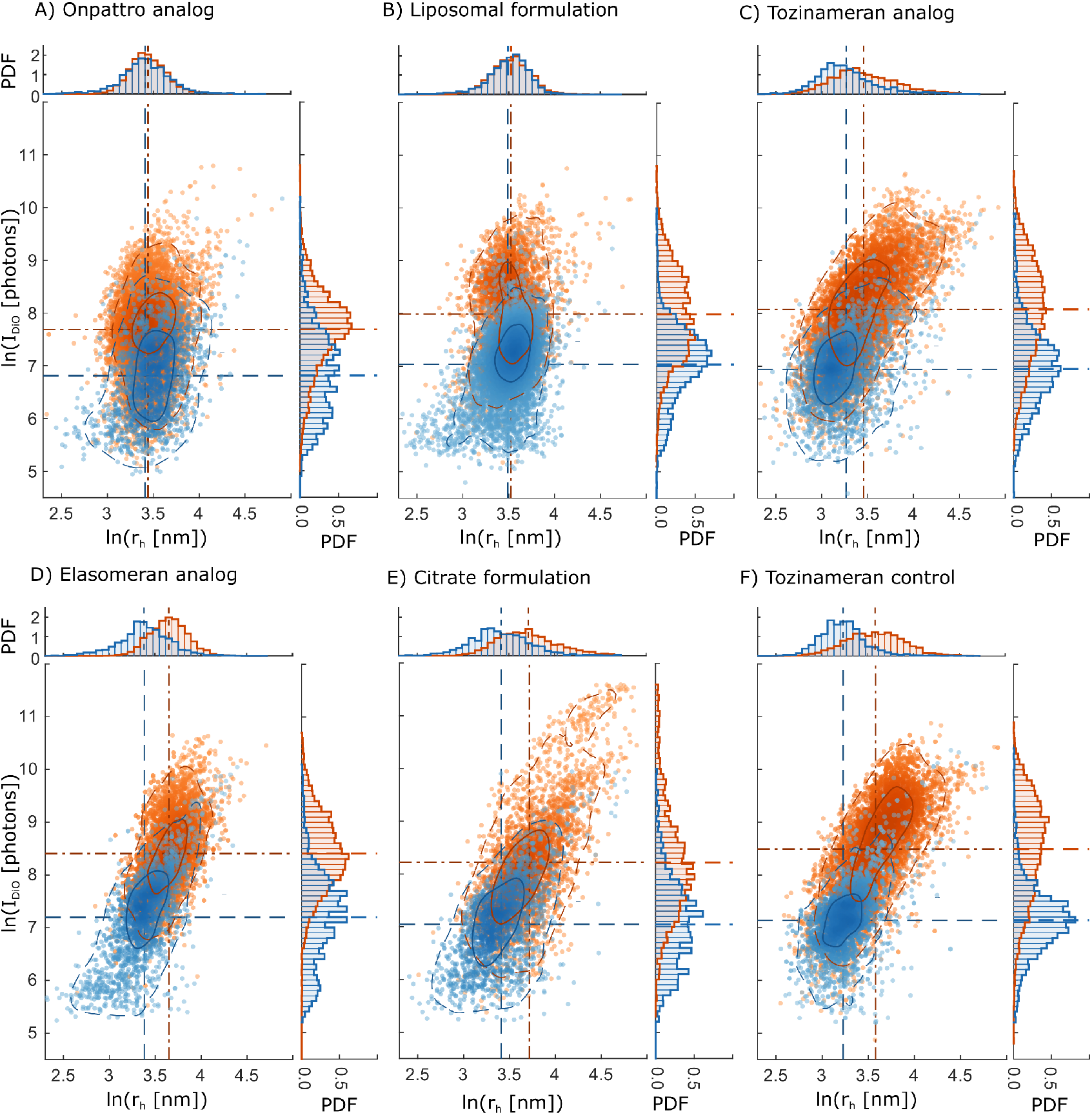
Formulation-dependent clustering of LNP subpopulations: Particle (DiO) intensities are plotted as a function of corresponding particle sizes, for subpopulations of loaded and empty LNPs, where individual particles are classified as loaded (containing Cy5-mRNA) or empty by looking for the presence Cy5 signal co-localized with DiO (lipid dye) signal. Results show that the Onpattro analog (A) and the Liposomal LNPs (B) have similar size distributions for their loaded and empty particles, while their corresponding intensities substantially increase for loaded LNPs. In contrast the Tozinameran analog (C), Elasomeran analog (D), Citrate (E) and the Tozinameran control (F) formulations show both increased particle sizes and intensities for their loaded subpopulations. The contour lines mark particle density, including a central region containing 68% (solid) of all particles and a wider region containing 95% (dashed) of all particles in each subpopulation.

We classify the formulations by calculating the differences in mean values of loaded and empty subpopulations, normalized by the pooled standard deviation – a statistical metric known as Cohen’s *d* (Methods, “Comparison of loaded and empty LNPs”). Our measurements reveal substantial separation in the DiO intensities of loaded versus empty particles, with *d* ≥ 0.5 across all formulations (Table 2). In contrast, the differences in particle size between loaded and empty LNPs vary considerably (Table 2). Hence, we categorize the formulations into three separate classes based on the magnitude of size differences between their subpopulations: 1) ‘Negligible-separation’ (*d* ≤ 0.2), observed for the Onpattro analog (Figure 2A) and the Liposomal LNPs (Figure 2B); 2) ‘Moderate-separation’ (0.2 *< d* ≤ 0.5), observed for the Tozinameran analog formulation (Figure 2C); and 3) ‘Substantial-separation’ (*d >* 0.5), observed for the Elasomeran analog, Citrate, and Tozinameran control formulations (Figures 2D, E & F).

**Table 2:**
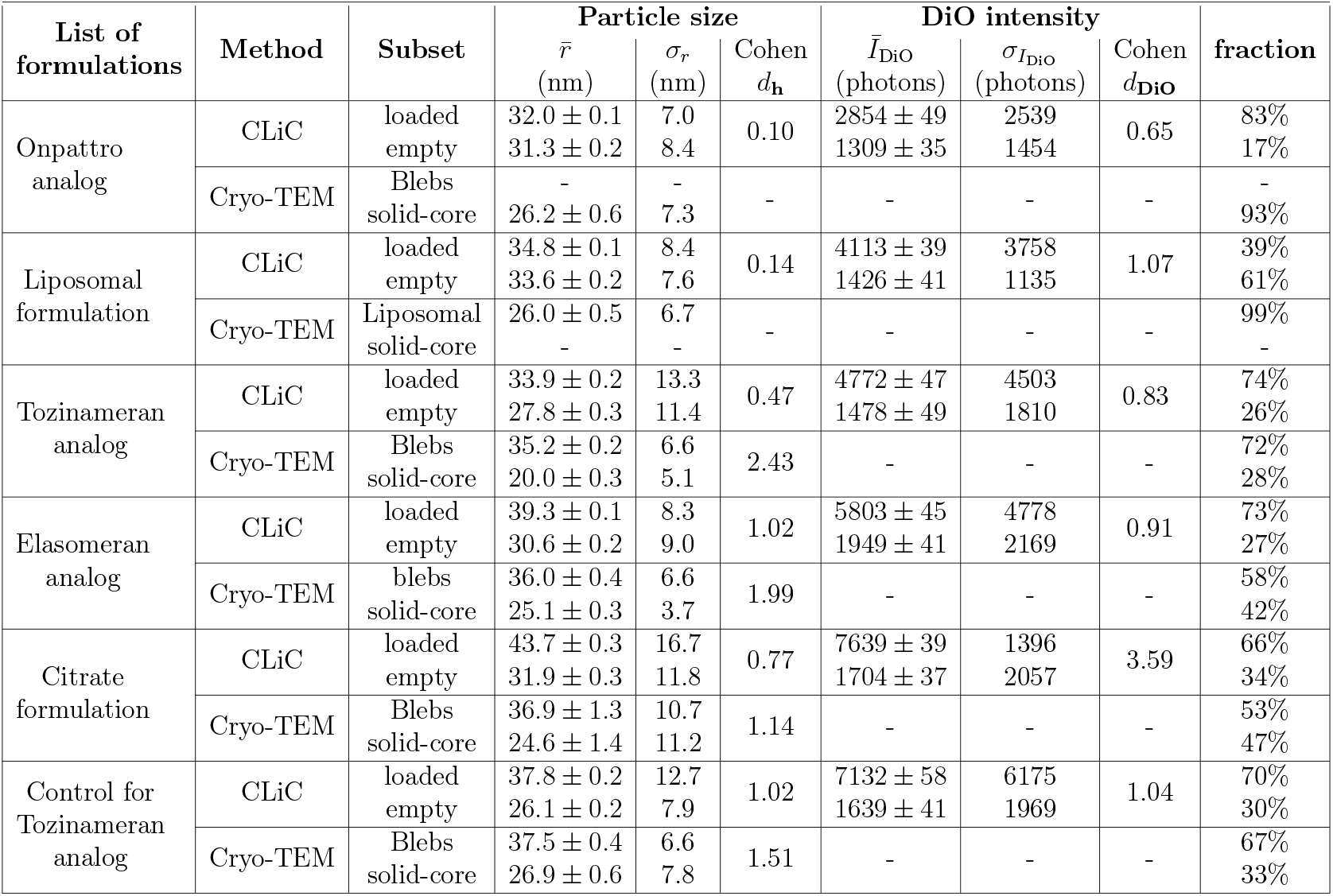
Summary of CLiC and Cryo-TEM measurements as a function of formulation parameters: CLiC measurements include mean size and DiO-intensity, where hydrodynamic radius (*r*_h_) is obtained from particle tracking. For Cryo-TEM measurements, the particle sizes are derived from projected areas of particles in TEM images, assuming spherical structures. The table compares size distributions of various subpopulations in each formulation; loaded versus empty particles for CLiC, and changes in particle structures for Cryo-TEM. Mean LNP radii 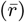, standard deviations (*σ*_*r*_), and Cohen’s d (*d*_**h**_, see Methods, “Comparison of loaded and empty LNPs”) values are compared for both techniques. Similarly, DiO intensities are compared through their mean 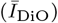, standard deviation 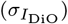 and Cohen’s d (*d*_**DiO**_), for CLiC measurements. Additionally, the table lists percentages of most LNP subpopulation presented. In cases of low statistics; where fewer than 50 particles were observed, the boxes are marked by dashes to indicate nonapplicable values. We observe that CLiC and Cryo-TEM measurements are consistent across all measurements; formulations that showed similar LNP sizes for loaded and empty subpopulations in CLiC also showed a single LNP morphology in Cryo-TEM measurements, while those that showed substantial changes in their particle size distributions showed two structures (*e*.*g*., LNP bleb versus nonbleb structures).

In the ‘negligible-separation’ class – namely the Onpattro and Liposomal formulations (Figure 2A, B) – we observe substantial differences in DiO signals (*d*_**DiO**_ *>* 0.6) despite their similar size distributions (*d*_**h**_ *<* 0.2). This implies that, at the single-particle level, loaded particles incorporate more lipid dyes, while their sizes are not substantially different from those of empty particles. Moreover, within each subpopulation we do not observe a clear correlation between particle size and DiO intensities, indicating that DiO incorporation, and hence per-particle lipid content, is heterogeneous.

In contrast, the ‘moderate-separation’ class – represented by the Tozinameran analog formulation (Figure 2C) – shows moderate differences in size distributions (*d*_**h**_ = 0.47) and more substantial differences in intensity distributions (*d*_**DiO**_ = 0.83) between loaded and empty subpopulations (Table 2). In addition, within each subpopulation, an increase in particle size correlates with an increase in DiO intensity.

Finally, in the ‘substantial-separation’ class – represented by the Elasomeran analog, Citrate and Tozinameran control formulations (Figures 2D, E & F) – we observe substantial differences in both size and DiO intensity for each of the loaded/empty subpopulations, and particle size correlates with DiO intensity within each subpopulation. Taken together, these in-solution single-particle measurements reveal a heterogeneous size and lipid-content distribution that define three structural particle classes, each with distinct loaded and empty subpopulations.

To further the investigation into LNP sample heterogeneity, we compare our CLiC-ALEX measurements to Cryo-TEM of particles with the same formulations to establish correlations between these methods (see Methods, “Cryo-TEM measurements” and Supplementary Information, Section 1.4). In the Cryo-TEM analysis, we classify subpopulations of particles based on the observed structures in images, where the presence of bleb structures or liposomal morphology indicates a high water content, in contrast to particles with solid-core structures.

In the ‘negligible-separation’ class, LNPs appear structurally homogeneous: the Onpattro analog formulation predominantly contains spherical LNPs with oil cores, while the Liposomal formulation consists mainly of liposome-like particles, as expected. In contrast, formulations in both the ‘moderate-separation’ and ‘substantial separation’ classes contain at least two subpopulations: one fraction of LNPs with bleb structures (Table 2) and another with solid-core structures, similar to those of the Onpattro analog. Interestingly, the relative fractions of LNPs identified by Cryo-TEM as either bleb-positive or bleb-negative are similar to those measured by CLiC-ALEX as either mRNA-positive or mRNA-negative. This similarity is most notable for the Tozinameran analog and Tozinameran control formulations, where the difference in fractions is within 3%. In contrast, for the Elasomeran analog and the Citrate formulations, the fractions of bleb-positive and mRNA-positive LNPs differ by ∼ 13 − 15%. Moreover, the difference in radius between bleb-positive and bleb-negative LNPs observed by Cryo-TEM is similar to the difference in radius between loaded and empty particles measured by CLiC-ALEX.

Overall, our results indicate that LNPs identified as having blebs in Cryo-TEM likely contain mRNA. However, depending on the formulation, a lack of bleb in a Cryo-TEM image is not necessarily an indication that the LNP does not contain mRNA, as shown by our quantitative CLiC-ALEX analysis. Moreover, while detecting a clear size difference between loaded and empty LNPs can be an indication of the presence of blebs in a sample, this correlation is also not strict.

### Distributions of mRNA copy number per LNP reveal formulation-dependent heterogeneity

Next, we compare our measured distribution of mRNA-number per LNP as a function of the formulation parameters, with our estimate of mRNA-number based on the LNP size, formulation, and N/P ratio, described in the Methods Section, “mRNA loading biophysical estimate”.

At the ensemble level, the measured mRNA distributions of loaded LNPs (orange histograms) show relatively good agreement with a biophysical estimate of the mRNA copy number (black histograms) for most formulations (Figure 3), with noted deviations for the Onpattro and Elasomeran analogs. The estimated LNP copy numbers are based on the LNP formulations’ measured particle size distributions (or equivalently, particle volume), and assumptions of, for example, the water fraction inside the LNPs (Methods, “mRNA loading biophysical estimate”). Thus, the difference in the estimated number of mRNA molecules per LNP 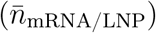 across formulations represents differences in LNP size distributions. In particular, deviations from the two histograms could indicate a different LNP water fraction or selective material losses during fabrication for these samples.

**Figure 3:**
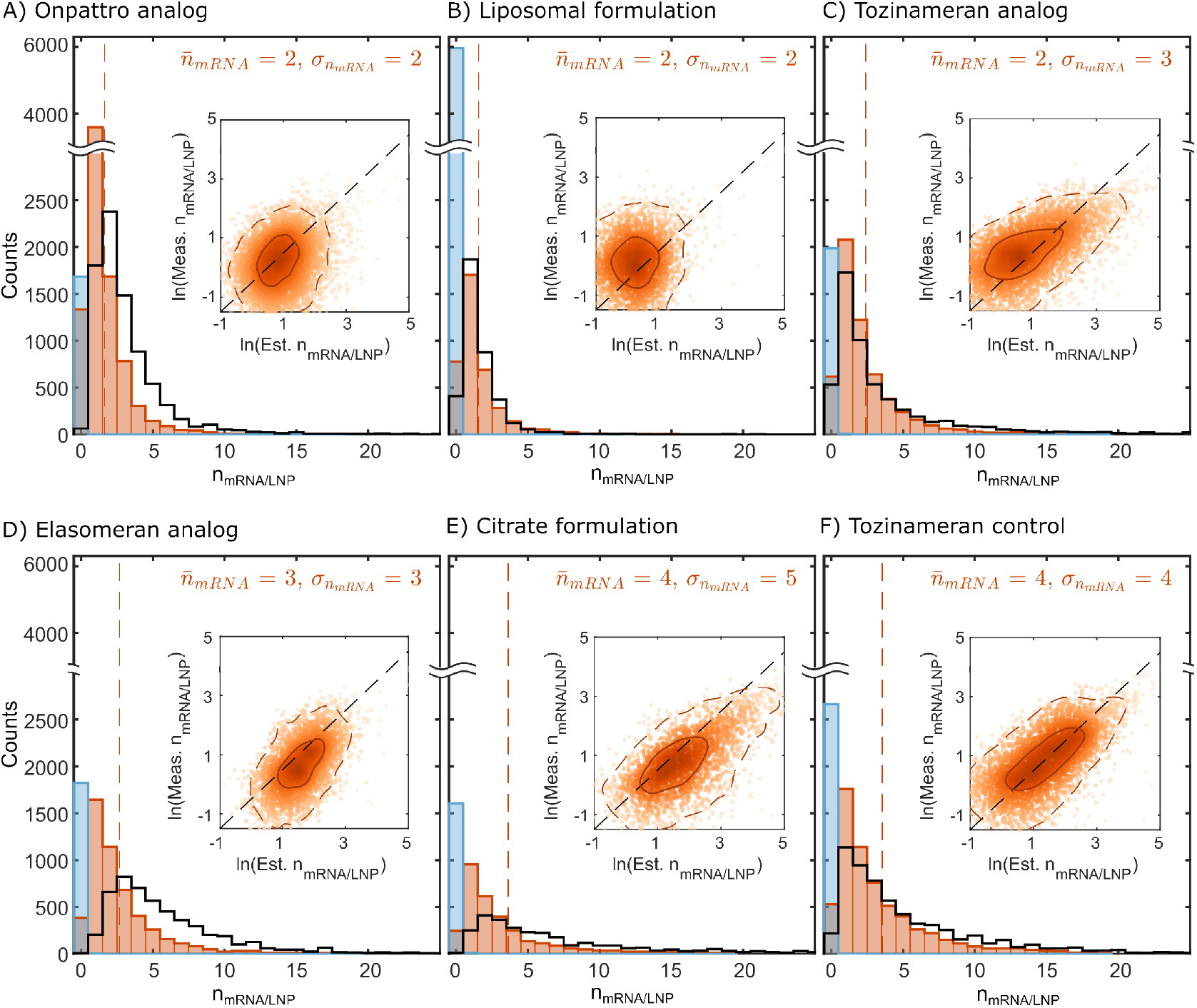
mRNA copy number per LNP varies by formulation. Single-particle measurements of Cy5 intensities per LNP were rescaled with the average single mRNA intensities. The resulting distributions of loaded LNPs (orange histograms) are in agreement with the expected single particle loading (black-line histogram) using a biophysical estimate (see Methods, “mRNA loading estimate”). Empty LNPs show low intensities in the Cy5 channel, consistent with no Cy5-mRNA cargo (blue histograms). Orange dashed lines indicate the mean values of corresponding distributions, which are also added as text labels. Scatter plots of measured mRNA signal versus the biophysical mRNA-loading estimate show two trends across LNP formulations, consistent with Figure 2. A) Onpattro analog and B) Liposomal LNPs show minimal scaling between mRNA content and the biophysical mRNA-loading estimate (particle volume), while the rest of the formulations (C, D, E and F) show positive scaling between the two parameters. The scatter plots also include 2-level density contour lines, highlighting regions with 68% (solid line) and 95% (dashed line) of all loaded particles, where the solid line represents regions with the highest density.

When investigating the scaling between particle size and measured mRNA copy number at the single-particle level (inset figures), we observe a varying degree of scaling for the different formulations. Here, formulations in the ‘negligible separation’ class – the Onpattro analog (Figure 3A insert) and the Liposomal LNPs (Figure 3B insert) – display a very weak scaling of the measured mRNA copy numbers per particle using CLiC-ALEX microscopy, versus the biophysical estimate for the mRNA copy number.

In contrast, the rest of the formulations showed relatively higher degrees of positive scaling, as shown by the alignment of the density contour lines with the reference dashed line (with a slope equal to one) across these formulations (inserts of Figure 3 C, D, E and F). Our results therefore indicate that different formulations and treatments of nanoparticle complexes can result in different degrees of heterogeneity in the mRNA loading, which may be due to steric and structural constraints imposed on the mRNA loading that are not fully taken into account by the biophysical estimate.

### Single-particle FRET measurements provide molecular arrangement information

To further investigate the structural properties of the mRNA-loaded LNPs, we compare CLiC-ALEX single-particle FRET measurements with particle structures obtained from Cryo-TEM images (Figure 4). For this, we plot the measured FRET efficiencies (*E*_FRET_) of individual particles against the corresponding FRET stoichiometry (*S*_FRET_, Methods, “Fluorescence signal quantification”).

**Figure 4:**
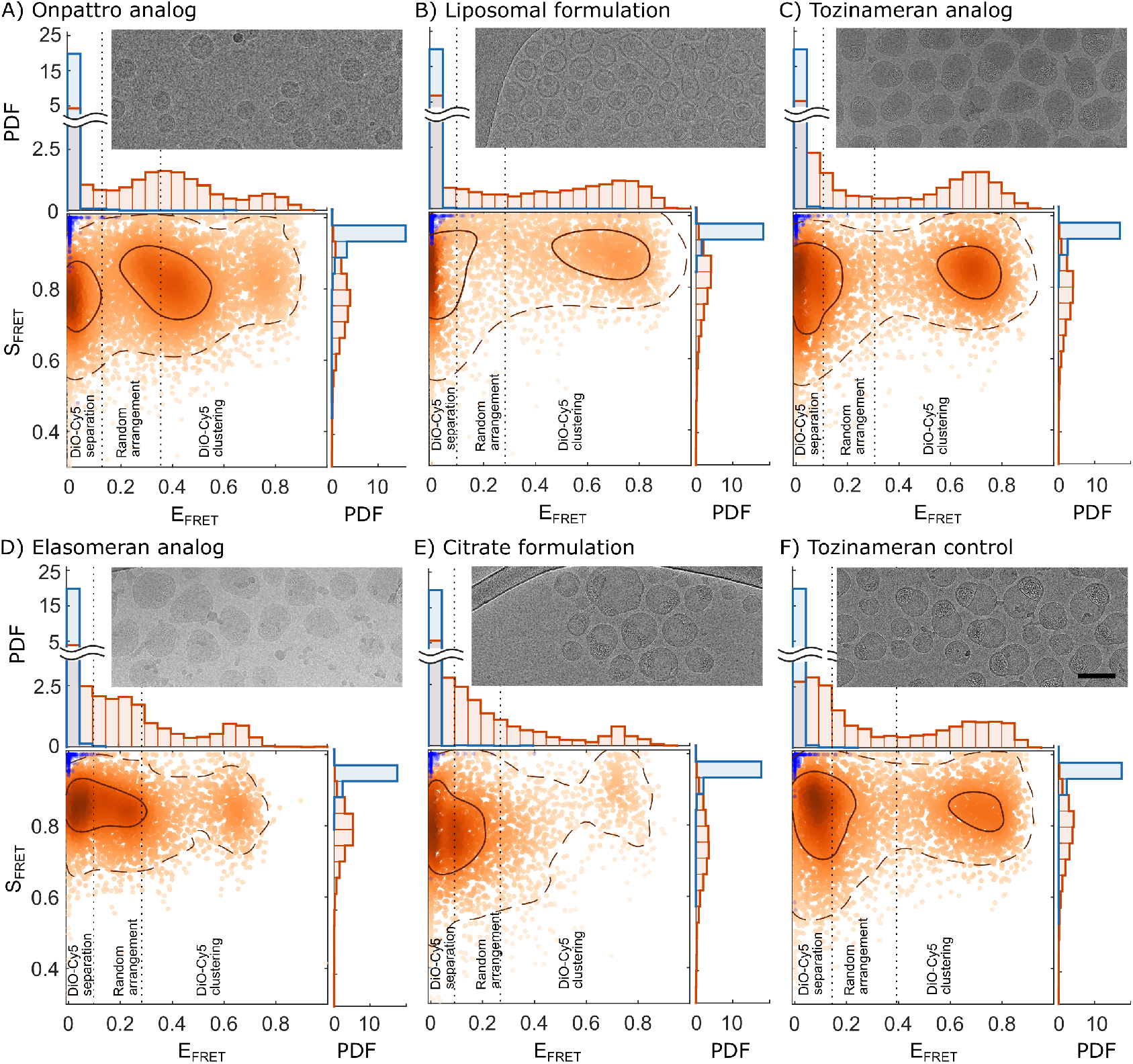
Measurements of FRET efficiencies and stoichiometry show distinct clustering as a function of formulation parameters: The results show substantial clustering on both scatter plots and projected histograms, for all formulations measured, including: A) the Onpattro analog; B) the Liposomal formulation; C) the Tozinameran analog; D) the Elasomeran analog; E) the Citrate formulation; and F) the control for the Tozinameran analog. Loaded LNPs are shown in orange and empty ones in blue. The loaded LNPs are overlayed with 2 level contour lines a solid line for regions containing 68% of all particles (prominent clusters) and dashed lines for 95% (representing the broader distributions). The scatter plot also includes vertical theoretical (dashed) lines, to highlight regions where different dye-dye interactions are expected, as a function of the DiO-Cy5 pair Förster radius and average dye concentrations (see Methods, “Relating FRET to molecular arrangement”). Overall, the clusters map to regions with high FRET (DiO-Cy5 clustering), medium FRET (due to random arrangement of dyes in LNPs), and low FRET (due to substantial separation between dye populations). The figures also include Cryo-TEM imaged for each formulation, which show LNP structures corresponding to the measured FRET profiles. Scale bar = 100 nm.

Most formulations have multimodal distributions of *E*_FRET_ in their loaded LNPs, and a single peak in the corresponding stoichiometry histograms (Figure 4). A similar stoichiometry value implies a similar relative amount of mRNA and DiO for each *E*_FRET_ cluster. Thus, the difference in *E*_FRET_ per cluster is associated with different arrangements of mRNA and DiO within the LNPs. To relate the *E*_FRET_ values to different molecular arrangements, the measured *E*_FRET_ values were compared to the expected *E*_FRET_ assuming a random arrangement of the dye molecules (Methods, “Relating FRET to molecular arrangement”). These estimates were then used to classify three regions of the scatter plots defined by the average separation between the donor and acceptor: 1) ‘DiO-Cy5 separation’ (low FRET); 2) ‘Random arrangement’ (medium FRET); and 3) ‘DiO-Cy5 clustering’ (high FRET). Overall, the observed clusters are not aligned with the formulation classes presented above, however, they can be understood based on possible lipid/mRNA arrangements within subpopulations of particles, for each formulation.

The low *E*_FRET_ cluster that is present for all of the formulations is likely due to the fact that most of the DiO is on the outer surface of the LNPs. Since we observe a *E*_FRET_ cluster close to zero, which occurs independently of whether blebs are detected, a low FRET signal is not necessarily due to the existence of blebs. Rather, taking into account the Förster radius of the DiO-Cy5 pair of approximately 5.0 nm (see Methods, “Relating FRET to molecular arrangement”), we can interpret a substantially lower FRET transfer efficiency than that expected from random molecular arrangement to indicate that DiO is located mainly on the outer surface of the LNPs in this cluster.

Following this reasoning, we can interpret the higher *E*_FRET_ clusters in Figure 4 to indicate that a substantial amount of internalized DiO is clustered together with the mRNA. Since DiO dyes carry a positive charge,^26^ we note that there could be an attraction between these dyes and the mRNA molecules, in particular at neutral pH when the ionizable lipids are uncharged; however Debye screening is expected to suppress such effects and we observe a strong dependence of the clusters on formulation. The high *E*_FRET_ cluster is also present across all formulations, at varying degrees of prominence, indicating that the internalization of DiO is such that the relative proximity of DiO to Cy5-mRNA molecules is heterogeneous.

The Onpattro analog formulation is unique in that it shows clusters across the three regions, with the dominant subpopulation located in the ‘random-arrangement’ region, followed by a secondary subpopulation in the ‘DiO-Cy5 separation’ region. For the Liposomal, Tozinameran analog and Tozinameran control formulations – whose structural features of loaded LNPs were previously noted in Cryo-TEM to align with either liposomal or bleb-like morphologies respectively – we observe at least two prominent fractions of LNPs: one in the ‘DiO-Cy5 separation’ region and another in the ‘DiO-Cy5 clustering’ region. In contrast, the Elasomeran and Citrate formulations each show a single dominant cluster in the ‘DiO-Cy5 separation’ region that extends into the ‘random-arrangement’ region, together with a smaller secondary cluster in the ‘DiO-Cy5 clustering’ region.

We note that for bleb-positive LNPs, a high *E*_FRET_ could potentially occur either when the mRNA is in the bleb with a high fraction of DiO in the inner bleb leaflet or when the mRNA is not in the bleb and there is a high fraction of DiO inside the LNP. In future work, super-resolution optical techniques may be able to distinguish between these cases, ^9^ and for each particle identify size, mRNA loading, bleb/not bleb phenotype, and FRET-intensity signal to investigate the correlation between different LNP structure measurables.

## Conclusions

CLiC-ALEX measurements provide a high-throughput single-particle approach for resolving and quantifying the heterogeneity in clinically relevant mRNA-lipid nanoparticles. This is achieved by combining simultaneous and sequential imaging of suspended LNPs trapped in microwells, enabling quantification of LNP size, lipid dye fluorescence, mRNA loading, and FRET on the single-particle level. These measurements reveal formulation-dependent heterogeneity across these parameters and identify subpopulations such as loaded and empty LNPs. By comparing with Cryo-TEM, we further assess the occurrence of blebs and relate structural features to the heterogeneity observed in CLiC-ALEX data.

Across formulations, we observe that loaded and empty LNPs separate into distinct subpopulations, with differences that are more pronounced in their lipid dye intensities than in hydrodynamic sizes, the latter quantified from particle diffusion measurements. This suggests that the presence of mRNA influences lipid packing and dye incorporation, even when the particle sizes are similar. The classification of formulations into three ‘separation’ classes of the mRNA and lipid molecules within the particles, which we define as ‘negligible’, ‘moderate’, and ‘substantial’ separation (quantified by Cohen’s *d* in the Results section) identifies a structural heterogeneity between the empty and loaded LNPs and highlights that it is formulation dependent.

The structural differences observed with CLiC-ALEX are aligned with the structural features observed by Cryo-TEM, thus providing independent validation of our approach. Our studies include formulations which correspond to particles with homogeneous structures, as well as others with at least two morphologies (e.g., bleb-like structures and solid-core LNPs). The overall agreement between the subpopulations identified by CLiC-ALEX optical measurements and Cryo-TEM provides a direct link between nucleic acid encapsulation and particle morphology. However, not all formulations showed a one-to-one match between the CLiC-ALEX and Cryo-TEM classification, highlighting the need for complementary measurements to fully resolve LNP heterogeneity and provide deeper insights into mRNA lipid interactions.

Measurements of mRNA copy number per LNP showed agreement with the biophysical estimate on the ensemble level, however, single particle analysis revealed broader variability across formulations. For example, while analysis of some formulations did not show a correlation between the measured mRNA copy number with the biophysical estimate, the majority did. This demonstrates the value of single-particle measurements in uncovering relationships that could be obscured by ensemble averaging and deepens our understanding of particle heterogeneity and structural details which ultimately impact therapeutic efficacy and potency.

FRET measurements add a further dimension by probing lipid-cargo arrangement. CLiC-ALEX in combination with FRET measurements revealed subpopulations in which DiO was confined to the particle surface and others in which it was distributed within the LNP volume. Moreover, since FRET can alter the quantification of the Cy5 signal by up to an order of magnitude in single-molecule microscopy, our work shows that it is important to take into account these effects for quantitative results. Our results show that the FRET signal can be used to investigate differences in molecular packing of mRNA within the LNP that could influence the stability, release, and bio-distribution of these complexes during drug delivery.

Together, our results show that mRNA-LNP formulations partition into subpopulations with distinct structural, compositional, and organizational characteristics. Recognizing and quantifying these subpopulations is essential for connecting formulation parameters with therapeutic performance and optimizing next-generation nanomedicines. The CLiC-ALEX method provides a broadly applicable platform for analysis of molecular complexes in solution with structural heterogeneity, enabling high-throughput, quantitative, and quick measurements that can complement existing tools such as Cryo-TEM. As the field advances toward increasingly sophisticated lipid and RNA formulations, multiparametric single-particle tools will be critical for quantitative optimization of drug formulation design, which we suggest will be most powerful if performed back-to-back with livecell imaging of the same formulations.

## MATERIALS AND METHODS

### Preparation of the lipid nanoparticles

The lipids DSPC, ESM, PEG-DMG, and ALC0159 were purchased from Avanti Polar Lipids. The ionizable lipids MC3, norMC3, and ALC0315 were purchased through the Ciufolini lab at UBC. The ionizable lipid SM102 was purchased from Cayman Chemicals. Cholesterol (Chol) was purchased from Sigma-Aldrich. Lipophilic dye, DiO, was purchased from ThermoFisher Scientific. Cy5 labeled mRNA (firefly luciferase) was purchased from APExBio (EZ Cap™ Cy5 Firefly Luciferase mRNA (5-moUTP), catalog No. R1010).

All LNP mRNA formulations were performed at an amine-to-phosphate ratio (N/P) of 6. The formulations used in the present study were prepared as follows:

#### 1. The Onpattro analog, Elasomeran analog and the control for Tozinameran

Lipid components (ionizable lipid/Chol/DSPC/PEG-DMG/DiO) were dissolved in ethanol at a final concentration of 10 mM total lipid with a molar ratio of 50/38.5/10/1.5/1 mol fraction, respectively. The ethanolic lipid phase was then rapidly mixed with an aqueous buffer (pH 4 25 mM NaOAc) containing mRNA through, using a custom made T-junction mixer in the Cullis lab (UBC). A flow rate ratio of 3:1 aqueous:ethanol phase (v/v) and a total flow rate of 20 mL/min was used for all formulations. Immediately after rapid mixing, the particles were then dialyzed overnight against a >500-fold volume of pH 7.4 PBS buffer.

#### 2. The Tozinameran analog

The same sample preparation process was used here, however, the lipid components (ALC0315/Chol/DSPC/ALC0159/lipophilic dyes) were mixed at molar ratio of 46.3/42.7/9.4/1.6/1 mol fraction, respectively.

#### 3. The liposomal formulation

The same sample preparation process was used here, however, the lipid components (norMC3/Chol/ESM/PEG-DMG/lipophilic dyes) were mixed at molar ratio of 20/40/40/1.5/1 mol fraction, respectively.

#### 4. The citrate formulation

The same sample preparation process was used here, however, the aqueous formulation buffer employed was pH 4 300 mM Na-citrate.

### Quality control analysis of LNP samples

Lipid concentrations were measured using the Cholesterol E Total-Cholesterol assay (Wako Diagonistics). Bulk particle size measurements were determined using DLS with a Malvern Zetasizer Ultra (see Supplementary Information, Section 4 for results). Additionally, particle size and fluorescent intensities were measured using NTA with Malvern’s Nanosight Pro instrument (see Supplementary Information, Section 4 for results).

### CLiC flow-cell cleaning, surface treatment, and assembly

In this study, a flow-cell design similar to those previously described^18,20,27^ was used (see Figure 1A of Kamanzi et. al., ACS Nano 2024^20^ for example flow-cells). The device consists of two glass coverslips (Ted Pella, product no. 260452; 200 ± 10 µm thick, 25 × 25 mm), separated by a 30 µm thick double-sided adhesive spacer (Nitto Denko, product no. 5603). The bottom coverslip is microfabricated with a series of cylindrical wells (0.5 µm in height and 3 µm in diameter) that serve as traps for the nanoparticles (see schematic in Figure 1B (ii)).

Both glass surfaces are passivated with 5 kDa polyethylene glycol (PEG) layers through silane chemistry and a cloud point PEGylation technique, as previously reported,^18^ but with modifications. Briefly, the surfaces of the glass coverslips were cleaned by sonication in acetone for 15 min, followed by sonication in isopropyl alcohol (IPA) for 15 min, and then cleaned in piranha (at a 3:1 mixing ratio of sulfuric acid (H_2_SO_4_, CAS: 7664-93-9, Thermo Fisher Scientific) and hydrogen peroxide (H_2_O_2_, CAS: 7722-84-1, Thermo Fisher Scientific) for 45 min. This process removes any dust particles and/or organic residues from the surfaces. The surfaces were then lightly etched in 1M KOH for 30 min at room temperature, to break surface bonds and increase the density of OH groups on the surface. Next, APTES ((3-aminopropyl)triethoxysilane, CAS: 919-30-2, Thermo Fisher Scientific) was deposited in a vapor phase^18^ inside a vacuum desiccator at approximately 1.5 Torr for 30 min at room temperature, followed by an annealing step at 120 ^°^C for 20 min on a hot plate, resulting in primary amines covering the surfaces of the coverslips. The flow-cell layers were then exposed to a solution of Succinimidyl Valerate (SVA) functionalized methoxy polyethylene glycol valeric acid (5K mPEG-SVA, Laysan Bio) in a 0.9 M sodium sulphate (Na_2_SO_4_, CAS: 7757-82-6, Thermo Fisher Scientific) solution for 30 min, forming a PEGylated layer on the surfaces. The coverslips were then thoroughly rinsed with deionized (DI) water (at least three rinses) and dried with high purity nitrogen gas before assembly of the flow-cell. The assembled flow-cell is then sealed in a custom microfluidic chuck.

### Single-particle CLiC microscopy

Single-particle imaging experiments were performed using CLiC microscopy^27–29^ in combination with alternating laser excitation. The imaging system included: a Nikon Ti-E inverted microscope, a coherent Sapphire 488 nm 150 mW LP laser, a coherent OBis 647 nm 120 mW laser, an acousto-optic tunable filter (AOTF) and its controller (Gooch & Housego), a function generator (BK precision, 4055, 50 MHz dual channel), two Andor EMCCD cameras (iXon Ultra 897), and a set of mirrors and dichroic mirrors listed below.

Excitation beams from the two lasers (488 and 647 nm) were co-aligned, through a set of mirrors (Thorlabs BB1-E02 and BB2-E02) and a dichroic mirror (Thorlabs DMLP605R), and used to excite samples trapped in the CLiC imaging flow cell. The laser beams were modulated by the AOTF, which was triggered synchronously with the Andor cameras by the function generator. The laser beams were then sent into a Nikon TiE inverted microscope, through a 100X oil imaging objective (Nikon, Apochromat TIRF 100× oil immersion objective lens N.A. 1.49, W.D. 0.12 mm, F.O.V 22 mm).

As previously described, a microfluidic chuck was used to secure the flow cell to a microscope stage, positioned between the objective lens and the CLiC pusher lens (Figure 1A). The CLiC pusher lens was controlled by a piezo-nanopositioner through the CLiC controller. Throughout the experiments, the CLiC lens was cyclically lowered and raised to trap nanoparticles within the wells and refresh them with new particles from the surrounding solution. This enabled quantitative measurements on thousands of individual nanoparticles, essential for characterizing heterogeneous samples.

The resulting emission signals were then collected by the same imaging objective and sent to the corresponding imaging cameras, through a set of dichroic mirror and emission filters (Chroma TRF59906, 488/640nm, Laser Dual Band filter set). In addition, fluorescent signals from dual-labeled samples were separated by a long pass filter (Chroma AT610lp), and sent to the pair of iXon Ultra 897 EMCCD cameras, both with a camera pixel size of 16 µm. Here we refer to the camera in the long-pass path as the red channel, while the other camera is referred to as the green channel, as shown in Figure 1A & C.

### Data analysis

The data analysis pipeline is described in detail in Section 1 of the Supplementary Information. However, the main steps are outlined below:

### Fluorescence signal quantification

To quantify the DiO signal (*I*_*DiO*_), we measure the total fluorescence signal on both cameras, with only the 488 nm laser turned on:

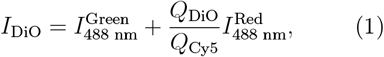

where the subscript for *I* indicates the illumination wavelength, the superscript for *I* indicates which emission channel is used, and *Q*_DiO_ and *Q*_Cy5_ are the quantum yields of DiO and Cy5, respectively, which are provided by the suppliers as 0.27 for Cy5 when in PBS and 0.51 for DiO.

To quantify the mRNA-Cy5 signal (*I*_*Cy*5_), we use a combination of simultaneous and sequential imaging:

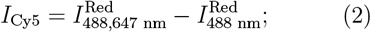

where 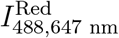 represents the red channel fluorescence with both lasers on, and 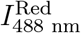 is the fluorescence in the same channel with only the blue laser one. This helps overcome the limitation of having to track weaker cargo signal and has the advantage of removing the potential contributions of both FRET and DiO bleedthrough to the Cy5 channel.

We quantify the FRET signal by estimating the FRET transfer efficiency:

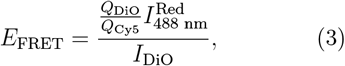

where *E*_FRET_ can be biophysically related to the spatial arrangement of DiO and Cy5-mRNA in LNPs (Supplementary Information, Section 1.4).

Stoichiometry (*S*_FRET_) is estimated as:

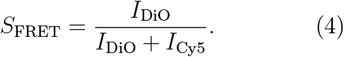

### Relating FRET to molecular arrangement

To relate *E*_FRET_ to molecular arrangement, we here approximate the dye distribution with 3D random arrangement, thus the approximation ignores the finite size of the LNPs. *E*_FRET_ in the case of random arrangement can be calculated from a known acceptor concentration:^30^

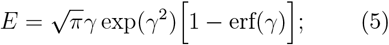

where:

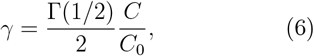

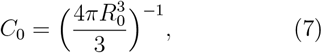

*R*_0_ is the FRET radius, and *C* is the Cy5 molecule concentration in the same length unit as the FRET radius. The Cy5 concentration *C* is here estimated for each formulation by taking the average measured particle volume of the loaded LNPs, while subtracting the contribution of the outer PEG layer, the measured mRNA copy number of loaded LNP, and the measured number of Cy5 labels per mRNA. The average particle volume and the measured mRNA copy number of loaded LNP comes from the CLiC-ALEX measurements. The measured number of Cy5 labels per mRNA was measured using Nanodrop One, a micro-volume spectrophotometer from Thermofisher Scientific, from which the average number of Cy5 labels per mRNA was measured to ≈ 15. *R*_0_ was estimated by comparing it with other FRET pairs with similar spectra that have a slightly higher spectral overlap with Cy5, such as Alexa488, as well as having a lower overlap, such as EGFP, while also compensating for the dependence on the donor quantum yield. Using this, the expected *R*_0_ for DiO-Cy5 is 5.0 ± 0.2 nm, where 5.0 is the mean value of the two and the upper/lower values are the FRET lengths for Alexa488/EGFP when compensating for the difference in quantum yields compared to DiO. The values were taken from https://www.fpbase.org/fret.

In the FRET plots there is a range corresponding to ±50% of *γ* to describe the uncertainty in the random arrangement estimate. That value is an estimate of the combined uncertainty in the estimate of the uncertain FRET radius and the uncertainties in the estimation of the Cy5 concentration on the single-particle level. Thus, any larger deviation than 50% of *γ* is here interpreted as an indication of either separation or clustering between the dyes. mRNA loading biophysical estimate:

The measured mRNA loading is here estimated by normalizing the measured LNP Cy5 signal with the Cy5 signal from free mRNA measured at the same conditions (Supplementary Information, Figure S3).

To compare the measured loading with the expected loading based on LNP size and molecular formulation we used the equation:

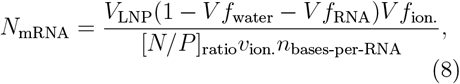

where *V*_LNP_ is the volume of the LNP, *V f* is the volume fraction for that component, [*N/P*]_ratio_ is the N/P ratio of the formulation, *v*_ion._ is the volume of individual ionizable lipids, and *n*_bases-per-RNA_ is the number of bases per mRNA molecule. Estimates of all of these factors except for *V*_LNP_ can be done using the molecular formulation, where those values are estimated in Supplementary Information, Section 1.3. Thus, the expected single-particle mRNA loading is estimated by those factors combined with a particle volume estimate from the measured particle radius assuming a spherical particle.

### Comparison of loaded and empty LNPs

Cohen’s d, used, to compare the differences between loaded and empty LNPs, is given by:

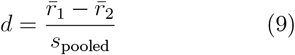

where *s*_pooled_ is given by:

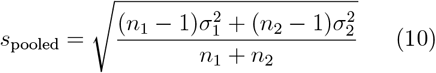

for two sample populations with 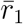 and 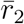 mean values, *σ*_1_ and *σ*_2_ standard deviation values, and *n*_1_ and *n*_2_ sample sizes.

### Colocalization classification

To improve the signal-to-noise (SNR) ratio during fluorescence signal estimate and colocalization classification, we here use position-assisted averaging.^16^ Here the particles are first tracked using DiO signal, which has high to crop particle-centered images in the Cy5 channel, which yields increased sensitivity in co-localizing weak cargo signals that may otherwise fall below our detection threshold (Supplementary information, Section 1.2). The particles are classified as containing cargo if the mRNA signal during simultaneous imaging is higher than an expected cross channel bleed-through signal ≤ 2%, where that value was estimated from measuring LNPs with DiO but without Cy5-mRNA (Supplementary Information, Figure S4).

### CLiC particle sizing

The mean squared displacement of a particle confined by a circle is described by:^18^

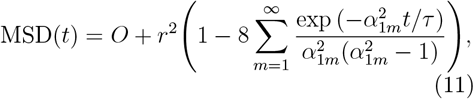

where *O* is an offset from position localization uncertainty, *r* is the confinement radius, *τ* = *r*^2^*/D* is the characteristic time, *D* is particle diffusivity, and *α*_1*m*_ is the *m*^th^ positive root of the derivative of the Bessel function of the first kind. To estimate the diffusivity in this work, the first two terms in the expansion are used. From that fit, the 2D diffusivity of the particle inside the well is extracted.

To convert diffusivity to size distributions, due to confinement and the proximity to nearby surfaces, we use a modified Stokes–Einstein relation:^25^

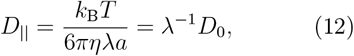

where *D*_||_ is the diffusivity of particles near two parallel confining planes, *D*_0_ is the diffusivity in bulk, *k*_B_ is the Boltzmann constant, *T* is the temperature, *η* is the kinetic viscosity of the solution, and *a* is the hydrodynamic radius. *λ* is a correction factor used to account for the hydrodynamic effects near surfaces and can be approximated as:^25,31^

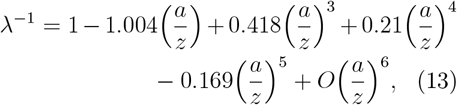

where *z* is the midway distance between the two planes of confinement. Given the known well depths (500 nm), the confinement effect is corrected by determining the correction factor that is self-consistent with the hydrodynamic diameter estimated from the Stokes-Einstein relation.^16^

### Cryo-TEM measurements

LNPs were concentrated to a lipid concentration of 15 - 20 mg/mL and 3 µL were added to glow-discharged copper grids and plunge-frozen in liquid ethane using a FEI Mark IV Vitrobot. Imaging was performed using a FEI LaB6 G2 TEM operating at 200 kV low-dose conditions paired with an FEI Eagle 4K CCD camera. Samples were imaged at a magnification of 55,000X. Cryo-TEM imaging was performed by the High Resolution Macromolecular Cryo-Electron Microscopy facility at the University of British Columbia (UBC).

The Cryo-TEM analysis was based on manual annotation, in which both the particle outline and bleb(s) were marked in MATLAB, as shown in Supplementary Information (Section 1.4). Particle sizes were subsequently estimated by relating the projected LNP area to a radius, with the assumption of an approximately spherical morphology.

## Supporting information

Data analysis pipeline and control measurements

## Associated Content

### Supplementary information

Supplementary information is provided, with the following topics:

- Detailed data analysis pipeline
- Control size measurements

## Acknowledgement

We acknowledge the substantial financial and other support from several granting agencies, including the Canadian Foundation for Innovation, British Columbia Knowledge Development Fund, PacifiCan and Western Diversification Fund, National Science and Engineering Research Council of Canada, Nanomedicines Innovation Network, as well as Startup Funds from the University of British Columbia Faculty of Science, Department of Physics and Astronomy, and Michael Smith Labs. Moreover, the Mitacs granting agency and industry sponsor ScopeSys co-supported the Postdoctoral fellowship held by A.K.. Further, E.O. held an international Postdoctoral fellowship from the Swedish Research Council (grant number 2024-00439) and Y.Z. held a NanoMedicines Innovation Network graduate award and a Canadian Institutes of Health Research Doctoral Award (FBD 193487). S.L. was supported by a Killam Accelerator Research Fellowship CryoTEM grid preparation and data collection was performed at the High Resolution Macromolecular Electron Microscopy (HRMEM) facility at the University of British Columbia (https://cryoem.med.ubc.ca). We thank Claire Atkinson, Amy Wo, Barathy Deivanayaga, Liam Worrall and Natalie Strynadka. HRMEM is funded by the Canadian Foundation for Innovation and the British Columbia Knowledge Development Fund.

## Author contributions

A. Kamanzi and S. Leslie contributed to the conceptualization and experimental design of the project. A. Kamanzi, A.T. Bido, and M. Stibbards-Lyle performed CLiC experiments used in the publication. Y. Zhang fabricated the LNPs with the assistance of J. Leung, M.H.Y. Cheng, and P.R. Cullis. A.T. Bido and E. Olsén performed DLS and NTA control measurements. E. Olsén performed the theoretical estimations and the Cryo-TEM analysis. A. Kamanzi, E. Olsén, and M. Jasinski developed the image analysis code with the assistance of B. Wang, M. Venier-Karzis and R. Berti. A. Kamanzi and E. Olsén performed the data analysis. A. Kamanzi and Y. Gu performed early proof-of-concept experiments, and C. Shaheen assisted with analyzing them. M. Jeliazkova and A. Kamanzi assisted with the micro-fabrication of the flow cells. S. Leslie supervised the project. A. Kamanzi, E. Olsén, and S. Leslie wrote the manuscript. All authors participated in discussion of the results and editing the manuscript.

## TOC Graphic

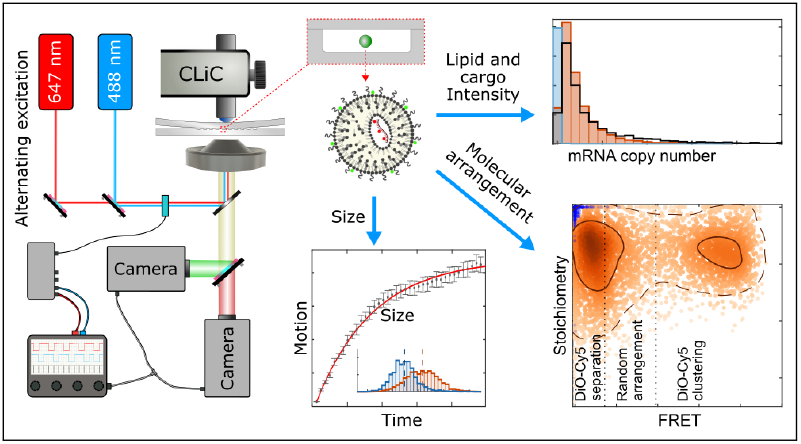

