## Supplementary material for "Single-particle multi-parametric microscopy reveals structural, size, and payload heterogeneity in mRNA-loaded lipid nanoparticles": Data analysis pipeline and control measurements

**References** **20**

### 1 Data analysis pipeline

#### 1.1 CLiC image analysis

As previously reported,[1] Convex Lens-induced Confinement (CLiC) measurements allow for single-particle tracking with long trajectories, where we here introduce a staggered laser excitation profile to achieve simultaneous and sequential dual-channel imaging conditions in a single measurement, as demonstrated by the image graphics in Figure 1C(ii) in the main text.

The image analysis pipeline is based on in-house scripts in MATLAB and Python and consists of multiple steps with different purposes:

1. Register wells using a brightfield image.
2. Detect wells containing a single diffusing particle.
3. For each well with a single diffusing LNP, denoise the image, track, quantify signals, and check co-localization.
4. For each trace, remove segments where the particle is temporarily stuck and, if the remaining track length is good enough, estimate the diffusivity.
5. If the trace has a low enough position uncertainty and a fitted well radius close to the physical well size, include the particle in the analysis.

###### 1.1.1 Well registration

Micro-well positions were detected using the OpenCV library in Python. First, brightfield images were collected in each experiment and used to identify the positions of micro-well centers and their radii. The initial candidate wells were then identified by computing a Hough transform of the brightfield images. This was implemented using the OpenCV's HoughCircles function, looping through a range of possible radii values – based on the know physical dimensions of the micro-wells – until a minimum threshold of candidate wells were detected.

Once candidate wells were identified, a grid template was constructed by computing the average radius of the wells and the pairwise distances between candidate wells to estimate the average horizontal and vertical spacing. Square regions around each well were summed and averaged over to compute a well template, and a Hough transform was calculated on the resulting average image to determine the radius of the averaged well.

Finally, a two-dimensional normalized cross-correlation was used to find areas in the white image with high correlation to the computed well template, providing more accurate locations of the wells. A radon transform was performed to determine the angles of the rows and columns of the wells.

##### 1.1.2 Detection of wells containing a single diffusing particle

The identification of wells containing a single diffusing particle was performed using a custom U-Net denoising (Section 1.2) and the count of the number of detections per well. The denoising and initial particle search was implemented in Python, and further filtration was performed to remove particles that stick to the surfaces of the wells was performed in MATLAB.

For each well position identified during grid registration, the system extracts a region of interest around the detected pit radius. The cropped regions undergo mask-based filtering to remove partial wells from adjacent positions to ensure that particle detection occurs only within the intended well boundaries. The regions were then denoised using a pre-trained U-Net to remove noise artifacts while preserving particle signals (Section 1.2). The initial detection of candidates used the Crocker-Grier method[2] implemented in the TrackPy library batch processing function to identify particle centers in each frame. The detection parameters include particle diameter estimates and minimum thresholds to distinguish particles from noise. The tracking is restricted to track the brightest candidate in each frame. Signal thresholding and overlap analysis were used to determine the number of particles present in each frame. The method utilizes configurable overlap percentages to handle cases where particles appear in close proximity. Wells are classified as containing single particles when the median count across all frames is equal to one. For wells that meet the single-particle criterion, motion analysis is performed to differentiate between freely diffusing and immobile (stuck) particles. The mean squared displacement (MSD) is calculated by evaluating the particle's positional displacements across multiple time lags. @ To validate the particle selection, manual selection was used as a benchmark. More than 16 thousand wells were analyzed, with over 800 containing particles, both by humans and by the Python pipeline. Human reviewers identified wells that met five key criteria: full confinement particle, presence of a single LNP, consistent free diffusion, no video cutoff, and sufficient signal-to-noise ratio. Bayesian analysis of the obtained data showed that the automated pipeline achieved >98% agreement with manual selections, with most discrepancies arising from human error or edge cases like low SNR or particle aggregation. This high concordance supports the reliability of the automated selection for identifying usable wells for downstream analysis.

###### 1.1.3 Denoising, tracking, and signal estimation

Denoising, tracking, and signal estimation were performed using MATLAB. For each well, all images for each laser combination are first denoised using a U-net1.2. When the 488 nm laser is on (DiO excitation), the LNPs are tracked using the DiO signal, whereas when only the 647 nm laser is on, the Cy5 signal is used for tracking. Each well is analyzed separately on the basis of the prior well identification and selection. Since the particles here only contain one particle, the particle tracking is based on maxima detection, where Gaussian fitting is used to achieve subpixel localization accuracy.

For signal quantification and colocalization, each laser combination is handled separately. For each particle detection, a small position-centered region of interest is saved in each imaging channel. If the particles are co-localized, there should be particles present in the center of each region of interest in both imaging channels, where the stack of images can be averaged to improve the signal-to-noise ratio when colocalizing weak particle signals. Due to bleaching, the max signal projection of the first 3 images is used for the signal estimation, where a particle is classified as colocalized when 1) there is a particle detection in both max projections which is less than 3 pixels apart, 2) the Gaussian fit in the cargo signal has an R-square value larger than 0.5, and 3) the cargo signal is higher than 2% of the DiO signal, as there is a weak bleed-through between the imaging channels (Figure S5). The Gaussian fits of the max projections are used to estimate both the DiO signal and the Cy5 signal.

##### 1.1.4 Trace evaluation

The particle trace is evaluated to minimize the risk of including noise traces while also ensuring that the traces correspond to a freely diffusing particle. First, only particle detections with an integrated signal larger than 10 is included, where the value comes from the values observed in noise detections. Second, since particles can be stuck for either the entire video or parts of the video, the trace was checked for stuck particles. This was done by checking if the movement between consecutive frames was more than 2 pixels, and if the displacement was less than 2 pixels for more than 15 consecutive frames, the interval of less than 2 pixels of motion was identified. If the start and end point of this interval were closer than 4 pixels, that whole frame interval was classified as stuck, which was then excluded from diffusivity estimation. Similarly, if a step between frames was greater than 5 times the median step length, that detection was classified as a noise detection. After these checks, if there were at least 120 particle detections when the particle is freely diffusing, they were passed along for the diffusivity estimate. If the trace was shorter than 120, the particle trace was excluded.

##### 1.1.5 Size estimation from diffusivity estimate

When relating the particle trajectories to hydrodynamic diameter, both the finite size of the well and the proximity between the particle and the surfaces in the well need to be considered. Details of how this was performed for the used geometry can be found in Ref.[1] Briefly, each particle trajectory is first converted to a mean squared displacement (MSD) curve, and only particles with at least 120 detections are included in the analysis. The resulting MSD curve is then fitted using a model that describes diffusion in a circular well,[1] with an example shown in Figure 1C of the main text. The mean squared displacement of a particle confined by a circle is described by[1]

$$\mathrm{MSD}(t) = O + r^2\Big(1 - 8\sum_{m=1}^{\infty}\frac{\exp\left(-\alpha_{1m}^2 t/\tau\right)}{\alpha_{1m}^2(\alpha_{1m}^2 - 1)}\Big), \tag{1}$$

where $O$ is a offset from position localization uncertainty, $r$ is the confinement radius, $\tau = r^2/D$ is the characteristic time, $D$ is particle diffusivity, and $\alpha_{1m}$ is the $m^{\mathrm{th}}$ positive root of the derivative of the Bessel function of the first kind. To estimate the diffusivity in this work, the first two terms in the expansion are used. From that fit, the 2D diffusivity of the particle inside the well is extracted. However, due to confinement and the proximity to nearby surfaces, the diffusivity of the particle in the well is lower than the corresponding bulk diffusion constant.[3]

To convert diffusivity to size distributions, we use a modified Stokes–Einstein relation:[4]

$$D_{||} = \frac{k_{\mathrm{B}}T}{6\pi\eta\lambda a} = \lambda^{-1}D_0, \tag{2}$$

where $D_{||}$ is the diffusivity of particles near two parallel confining planes, $D_0$ is the diffusivity in bulk, $k_{\mathrm{B}}$ is the Boltzmann constant, $T$ is the temperature, $\eta$ is the kinetic viscosity of the solution, and $a$ is the hydrodynamic radius. $\lambda$ is a correction factor used to account for the hydrodynamic effects near surfaces and can be approximated as[4,5]

$$\lambda^{-1} = 1 - 1.004\Big(\frac{a}{z}\Big) + 0.418\Big(\frac{a}{z}\Big)^3 + 0.21\Big(\frac{a}{z}\Big)^4 - 0.169\Big(\frac{a}{z}\Big)^5 + O\Big(\frac{a}{z}\Big)^6, \tag{3}$$

where $z$ is the midway distance between the two planes of confinement. Given the known well depths (500 nm), the confinement effect is corrected by determining the correction factor that is self-consistent with the hydrodynamic diameter estimated from the Stokes-Einstein relation.[6]

#### 1.2 U-Net for fluorescence denoising

The simulations for the network training were performed using DeepTrack 2.0,[7] a Python-based software package developed for simulations of optical systems and training of neural networks. The network used is based on the U-Net architecture,[8] and its details are shown in Figure 1. Aside from the information given in the figure caption, the input is an image of size $(32 \times 32)$, there are three convolution steps per pooling step, and the network has in its up- and downsampling path $\{16, 32\}$ filters in the base block $\{32, 32\}$ filters and in the last step $\{32, 32\}$ filters. In total, the network contains 113,025 trainable parameters.

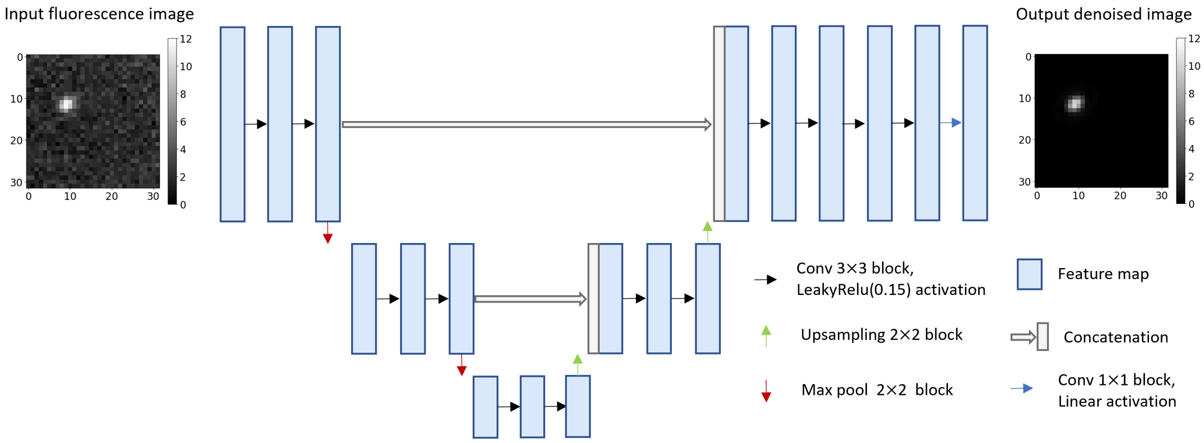

Figure S1: **U-net architecture**. The U-Net architecture comprises of four downsampling and upsampling blocks. The downsampling path has convolutional blocks with a 3x3 kernel, Relu activation, and 2x2 max pool downsampling. In the upsampling path, there are transposed convolutional blocks with a 3x3 kernel and Relu activation. Within the base block, convolutional blocks are executed, and a linear activation is applied in the final convolutional layer.

The input to the network is a fluorescence image of size $(32 \times 32)$, where the initial normalization of the image consists of subtracting the image with its minimum value and dividing the image with the standard deviation of the image. The network output consists of a particle image in focus, with minimized noise, corresponding to the input fluorescence image with all the noise and background signals removed. This output is then multiplied by the same standard deviation value used to normalize the input data to the U-Net. An example of this input-output pair is shown in the top left and right of Figure 1. The same network was used to denoize both Cy5-mRNA fluorescence and DiO fluorescence.

The training is based on the simulation of a fluorescence point particle where the particles are placed with ±2.0 µm from the center of the image and the depth position ranges from ±0.5 µm. The range of particle signals during training ranged from a peak signal of 5 to 10000, to span the range of signal values occurring experimentally, where the background noise is Poisson noise with a $\lambda$ ranging between 1-10. In addition to noise, the background signal consists of a small random linear gradient, as well as a weak signal from a ring-like structure with a random radius in the range of 2-3 µm, where the latter mimics the potential signal from the edge of the wells. Moreover, the numerical aperture and optical aberrations were chosen so that they mimic the experimental data. Furthermore, during the training, 90% of the images contain a particle and 10% are empty.

Training a neural network involves feeding data through a network, adjusting the weights to minimize the error via backpropagation, and repeating this process for multiple epochs. Our network receives data through a generator that generates new data samples for each batch, using the simulation pipeline above. For the training of the network, the mean absolute error (of the pixels) is used as the loss function,

$$\mathcal{L}_{\text{mae}} = \frac{1}{N} \sum_{i=1}^{N} |x_i - y_i|, \tag{4}$$

where $x_i$ represents the label and $y_i$ the prediction, which in this case are two images. Furthermore, the Adam optimizer[9] is chosen with a learning rate of $8e-5$, and $\beta_1 = 0.9$ and $\beta_2 = 0.999$. Training is carried out with a batch size of 32 until $L_{mae}$ no longer improves. Training is performed using the DeepTrack 2.0 package, with Python 3.10.10 and Tensorflow 2.10.0. The hardware configuration features an NVIDIA GeForce RTX 4070 GPU, an Intel Core i9 12900KF processor, 64 GB of memory, and runs on Windows.

#### 1.3 Parameter estimations when relating CLiC-ALEX data to expected mRNA loading

To estimate the expected number of mRNA copies inside an LNP (Eq. 8 in the main text), parameters such as the water content ($Vf_{\mathrm{water}}$), volume fraction of ionizable lipids ($Vf_{\mathrm{ion.}}$), and volume fraction of RNA ($Vf_{\mathrm{RNA}}$) need to be estimated. The water content in the core of an LNP is typically between 20-35 vol%[10–12] and the volume fraction of the mRNA cargo is often around 2-5 vol%.[10–12] Thus, given a certain variation in the water content and cargo loading, the volume fraction of the LNP containing lipids can be approximated with $(1 - Vf_{\mathrm{water}} - Vf_{\mathrm{RNA}}) = 0.65 \pm 0.2$. Furthermore, the volume fraction of the ionizable lipid $Vf_{\mathrm{ion.}}$ with respect to the total volume of lipids in the LNP can be estimated from the lipid molar ratio using that cholesterol has a volume around 0.63 $\mathrm{nm}^3$[13] whereas the other lipids in the LNP have an approximate volume of around 1.2 $\mathrm{nm}^3$. For the lipid molar ratio used, for all LNPs except long-circulating LNPs, $Vf_{\mathrm{ion.}} \approx 62\%$. Combined with the N/P ratio of 6 and the mRNA length of 1929 bases, the number of mRNA per LNP is

$$N_{\mathrm{mRNA}} = \frac{V_{\mathrm{LNP}}(1 - Vf_{\mathrm{water}} - Vf_{\mathrm{RNA}})Vf_{\mathrm{ion.}}}{[N/P]_{\mathrm{ratio}} v_{\mathrm{ion.}} n_{\mathrm{bases\ per\ RNA}}}, \tag{5}$$

where $V_{\mathrm{LNP}}$ is the volume of the LNP and $v_{\mathrm{ion.}}$ is the volume of individual ionizable lipids. Here, $V_{\mathrm{LNP}}$ can be estimated by the hydrodynamic radius by assuming a spherical LNP, while $v_{\mathrm{ion.}}$ values are obtained from literature. Thus, the expected mRNA loading from the biophysical estimation is a rescaling of the measured particle volume.

#### 1.4 Analysis of Cryo-TEM images:

We used cryo-TEM images to obtain complementary particle size information, as well as classify different particle structures observed as a function of formulation parameters. This included spherical non-bleb structures, bleb structures and liposomal LNP morphologies.

The cryo-TEM image analysis is here based on manual annotation, where the outline of

LNP as well as the bleb(s) was marked using the function "drawpolygon" in MATLAB. The particle size was estimated by calculating the area of the LNP, and then converting the area to a radius assuming a spherical particle. For each LNP, the number of blebs and their size was also saved, allowing estimation of the prevalence of LNPs containing blebs, as well as the LNP size for each particle subpopulation.

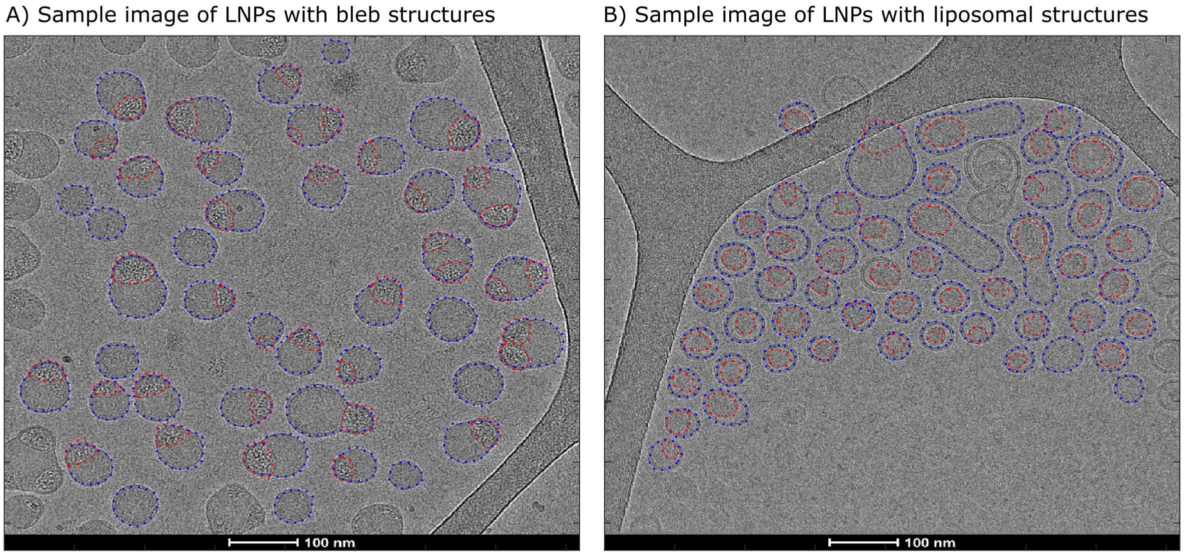

Figure S2: **Analysis of cryo-TEM images for classification of various LNP structures. (A-B)** cryo-TEM images as well as the manually annotated outline where the outline of the LNP is in blue and the outline of the bleb(s) or the solid core (for liposomal LNPs) is in red.

Figure S2 shows example images, including LNPs with bleb structures (Figure S2 A) and LNPs with liposomal particle structures (Figure S2 B). In Figure S2 A, each particle outline is marked with a blue dotted line, and particles that show phase separation of their lipid and aqueous volumes are also marked with red dotted lines around their aqueous volumes. In contrast, for liposomal LNPs (Figure S2) – which show an outer bilayer structure and an internal oil droplet – are marked with a red line around their lipid droplet. Using this analysis, we quantified the fraction of LNPs containing these structures by measuring the fraction of their areas made up of aqueous contents. The resulting plots are shown below in Figure S3.

### 2 Control measurements

#### 2.1 Free mRNA molecules

Reference measurements were performed on freely diffusing Cy5-labeled mRNA molecules to quantify the Cy5 signal per single mRNA molecule. The average intensity of a single mRNA molecule was then used to estimate the number of mRNA molecules in each lipid nanoparticle by normalizing the measured mRNA signal from the LNPs.

Figure S4 below shows the resulting distribution of Cy5 intensities for individual mRNA molecules. A total of 830 molecules were detected and analyzed, with a median intensity of 227 photons, with a mean of 263 photons, and standard deviation of 130 photons. Due to the non-zero width of the distribution, this implies that there is a corresponding uncertainty in the number loading on the single particle level, which affects the width of the obtained mRNA copy number distribution in Figure 4 in the main text.

Here, CLiC flow-cells were prepared by using piranha cleaning, as previously described.[14] However, surface PEGylation was not required as mRNA molecules do not bind to piranha-clean glass surfaces because of charge repulsion. This choice did not affect our imaging conditions, as there was no detectable difference in the background signals of plain glass vs those that were PEGylated.

The mRNA molecules were prepared by diluting them in 1X PBS buffer, to an imaging concentration of 0.1 nM, which allowed for single molecules per CLiC micro-well during imaging. All the other imaging conditions were kept similar to those used for LNP samples, with CLiC ALEX-FRET excitation settings used. This included laser powers of 6 mW for the 488 nm Coherent laser and 5 mW for the 647 nm laser (both measured at the objective), a camera exposure time of 6 ms and camera frame rate at 160 frames per second. The resulting images of freely diffusing mRNA molecules were then analyzed using sequential acquisition, as described for Figure 1(D i.) of the main text, where a frame mask was used to select frames in which the Cy5 dyes were excited. The only difference is that the free mRNA was tracked using the Cy5 signal, whereas the LNP in this work were tracked using the DiO signal.

#### 2.2 LNPs with unlabeled mRNA cargo

Figure S5 compares the Citrate formulation presented in the main paper with a control Citrate formulation containing unlabeled mRNA molecules. These measurements were performed in order to quantify cross-channel bleed-through of the DiO dyes into the Cy5 channel of our imaging system. Hence, they allowed us to determine the minimum threshold of the detectable Cy5 signal from loaded LNPs, where we expect a cross-channel bleed-through of the DiO signal to be less than 2% because of the used optical filters.

Figure S5 compares the simultaneous imaging of LNPs with unlabeled mRNA (Figure S5 A) versus LNPs with Cy5-labeled mRNA (Figure S5 D), where LNPs are labeled with DiO dyes. As expected, our measurements showed that less than 2% DiO fluorescence is detected in the Cy5 channel, for the formulation that contains unlabeled cargo. This is shown by a dashed black line, which represents this minimum detection threshold. The majority of the LNPs in Figure S5 have no fluorescence signal in the Cy5 channel, as shown by the linear cluster at the bottom of the figure. However, a smaller fraction of the particles show an increasing signal in this channel as the DiO signal increases. This can be attributed to cross channel bleed-through of the DiO signal, however, these intensities stay below the 2% threshold.

Furthermore, these measurements served as a check for the possible effects of using Cy5-labeled mRNA versus unlabeled one. This was verified by comparing the size and intensity distributions of both sets of formulations. Both Figures S5 A and D include projected histograms of DiO intensities, as well as the corresponding mean intensities (vertical dashed lines). Figure S5 D also includes a projected histogram of all particles in this formulation, and corresponding mean intensity (shown by the black histogram and vertical lines, respectively). When comparing the DiO intensity of the control LNPs (Figure S5 A, histogram) to those of the combined distribution of the Citrate formulation (Figure S5 A, black histogram), we observe that they have similar profiles, although those with Cy5-mRNA (Figure S5 A) show a broader distribution.

Figures S5 B and E show the total DiO fluorescence ($I_{\mathrm{DiO}}$, see Equation 1 of main paper) plotted as a function of the hydrodynamic radii ($R_{\mathrm{h}}$) of the particles. Note that Figure S5E is reproduced from Figure 2 of the main paper, with the addition of the combined loaded and unloaded LNP histograms (in black). Here, we observe that the average particle size ($\bar{R}_{\mathrm{h}}$) remains the same when using Cy5-labeled mRNA instead of unlabeled mRNA. This is shown by comparing the vertical blue line in Figure S5 B to the vertical black line in Figure S5 E, as well as the corresponding horizontal histograms. However, similar to the DiO measurements in Figure S5 A and D, we observe a broadening of the measured size distribution when using LNPs containing Cy5-mRNA. This is shown by the PDI of the two distributions, where LNP containing unlabeled mRNA had a PDI of 0.056 versus 0.145 for those with labeled mRNA cargo. Although this may indicate potential effects of Cy5 on the nanoparticles, additional measurements are necessary to further explore these observations.

Figures S5 C and F compare the total Cy5 fluorescence ($I_{\mathrm{Cy5}}$, see equation 2 of main paper) of the two formulations. Again we observe little to no Cy5 signal for most particles in the control formulation (Figure S5 C), with the exception of a few particles (133 particles out of a total of 2380). While the CLiC-ALEX assay is designed to minimize contributions from cross-talk and dye interactions, the observed signal in this small fraction of LNPs (less than 6% of all measured particles) can be explained by cross-talk in the system.

#### 2.3 LNP size measurements comparison: CLiC, cryo-TEM, DLS and NTA

Table 1 compares CLiC size measurements to cryo-TEM, dynamical light scattering (DLS) and nanoparticle tracking analysis (NTA). Here, complementary size distribution measurements using NTA were performed using Nanosight Pro from Malvern. The LNPs were diluted between $1 : 10^4 - 10^5$ in PBS water such that the concentration during the measurement was close to $10^9$ particles per ml, where the PBS water was first filtered through a 0.02 μm syringe filter (Whatman Anotrop, Cytiva) such that the particle count of the solution without particles was one or less per field of view.

Table 1: **Summary of CLiC, Cryo-TEM, DLS and NTA measurements as a function of formulation parameters:** CLiC measurements include size and DiO-intensity, while Cryo-TEM, DLS and NTA show the corresponding particle sizes. The table compares size distributions of loaded and empty LNPs, using their mean hydrodynamic radii ($\bar{r}_\mathrm{h}$), polydispersity index (PDI), standard deviations ($\sigma_{r_h}$), as well as the corresponding Cohen's d ($d_{r_\mathrm{h}}$, see SI). Similary, the DiO intensities are compared through their mean ($\bar{I}_\mathrm{DiO}$), standard deviation ($\sigma_{I_\mathrm{DiO}}$) and Cohen's d ($d_{I_\mathrm{DiO}}$). Additionally, the table lists the number of LNPs obtained for each subpopulation, as well as the percentage of loaded LNPs in each formulation.

| **List of formulations** | **Method** | **Subset** | **number of LNPs** | **Particle size** | | | | **DiO intensity** | | | **fraction** |
|---|---|---|---|---|---|---|---|---|---|---|---|
| | | | | $\bar{r}$ (nm) | $\sigma_r$ (nm) | PDI | Cohen $d_r$ | $\bar{I}_\mathrm{DiO}$ (photons) | $\sigma_{I_\mathrm{DiO}}$ (photons) | Cohen $d_{I_\mathrm{DiO}}$ | |
| Onpattro analog | CLiC | loaded | 8086 | $32.0 \pm 0.1$ | 7.0 | 0.048 | 0.10 | $2854 \pm 49$ | 2539 | 0.65 | 83% |
| | | empty | 1687 | $31.3 \pm 0.2$ | 8.4 | 0.073 | | $1309 \pm 35$ | 1454 | | 17% |
| | Cryo-TEM | Blebs | 10 | $35.2 \pm 2.5$ | 7.9 | 0.051 | n/a | n/a | n/a | n/a | 7% |
| | | Nonblebs | 130 | $26.2 \pm 0.6$ | 7.3 | 0.079 | | | | | 93% |
| | DLS | n/a | n/a | 22.4 | 9.2 | 0.171 | n/a | n/a | n/a | n/a | n/a |
| | NTA | n/a | n/a | 41.5 | 33.4 | 0.647 | n/a | n/a | n/a | n/a | n/a |
| Tozinameran analog | CLiC | loaded | 5688 | $33.9 \pm 0.2$ | 13.3 | 0.155 | 0.47 | $4772 \pm 47$ | 4503 | 0.83 | 74% |
| | | empty | 1995 | $27.8 \pm 0.3$ | 11.4 | 0.167 | | $1478 \pm 49$ | 1810 | | 26% |
| | Cryo-TEM | Blebs | 806 | $35.2 \pm 0.2$ | 6.6 | 0.036 | 2.43 | n/a | n/a | n/a | 72% |
| | | Nonblebs | 310 | $20.0 \pm 0.3$ | 5.1 | 0.066 | | | | | 28% |
| | DLS | n/a | n/a | 27.5 | 13.1 | 0.231 | n/a | n/a | n/a | n/a | n/a |
| | NTA | n/a | n/a | 43.8 | 27.6 | 0.397 | n/a | n/a | n/a | n/a | n/a |
| Elasomeran analog | CLiC | loaded | 5044 | $39.3 \pm 0.1$ | 8.3 | 0.045 | 1.02 | $5803 \pm 45$ | 4778 | 0.91 | 73% |
| | | empty | 1825 | $30.6 \pm 0.2$ | 9.0 | 0.086 | | $1949 \pm 41$ | 2169 | | 27% |
| | Cryo-TEM | blebs | 220 | $36.0 \pm 0.4$ | 6.6 | 0.033 | 1.99 | n/a | n/a | n/a | 58% |
| | | Nonblebs | 162 | $25.1 \pm 0.3$ | 3.7 | 0.022 | | | | | 42% |
| | DLS | n/a | n/a | 28.7 | 7.4 | 0.068 | n/a | n/a | n/a | n/a | n/a |
| | NTA | n/a | n/a | 36.8 | 14.7 | 0.159 | n/a | n/a | n/a | n/a | n/a |
| Citrate formulation | CLiC | loaded | 3155 | $43.7 \pm 0.3$ | 16.7 | 0.145 | 0.77 | $7639 \pm 39$ | 1396 | 3.59 | 66% |
| | | empty | 1612 | $31.9 \pm 0.3$ | 11.8 | 0.136 | | $1704 \pm 37$ | 2057 | | 34% |
| | Cryo-TEM | Blebs | 72 | $36.9 \pm 1.3$ | 10.7 | 0.084 | 1.14 | n/a | n/a | n/a | 53% |
| | | Nonblebs | 64 | $24.6 \pm 1.4$ | 11.2 | 0.21 | | | | | 47% |
| | DLS | n/a | n/a | 51.3 | 22.3 | 0.189 | n/a | n/a | n/a | n/a | n/a |
| | NTA | n/a | n/a | 50.3 | 29.6 | 0.347 | n/a | n/a | n/a | n/a | n/a |
| Liposomal formulation | CLiC | loaded | 3789 | $34.8 \pm 0.1$ | 8.4 | 0.059 | 0.14 | $4113 \pm 39$ | 3758 | 1.07 | 39% |
| | | empty | 5958 | $33.6 \pm 0.2$ | 7.6 | 0.051 | | $1426 \pm 41$ | 1135 | | 61% |
| | Cryo-TEM | Liposomal | 158 | $26.0 \pm 0.5$ | 6.7 | 0.67 | n/a | n/a | n/a | n/a | 99% |
| | | Nonliposomal | - | - | - | - | | | | | - |
| | DLS | n/a | n/a | 24.7 | 8.9 | 0.139 | n/a | n/a | n/a | n/a | n/a |
| | NTA | n/a | n/a | 38.5 | 31.8 | 0.684 | n/a | n/a | n/a | n/a | n/a |
| Control: Tozinameran analog | CLiC | loaded | 6283 | $37.8 \pm 0.2$ | 12.7 | 0.112 | 1.02 | $7132 \pm 58$ | 6175 | 1.04 | 70% |
| | | empty | 2667 | $26.1 \pm 0.2$ | 7.9 | 0.091 | | $1639 \pm 41$ | 1969 | | 30% |
| | Cryo-TEM | Blebs | 322 | $37.5 \pm 0.4$ | 6.6 | 0.031 | 1.51 | n/a | n/a | n/a | 67% |
| | | Nonblebs | 157 | $26.9 \pm 0.6$ | 7.8 | 0.083 | | | | | 33% |
| | DLS | n/a | n/a | 31.1 | 13.2 | 0.183 | n/a | n/a | n/a | n/a | n/a |
| | NTA | n/a | n/a | 40.5 | 18.5 | 0.208 | n/a | n/a | n/a | n/a | n/a |

The sample was illuminated using a 488 nm laser module, and for the fluorescence measurements a 500 nm long-pass filter was inserted. All measurements were performed under flow to increase statistics and minimize bleaching, and at least 10 videos of each image modality were recorded. The complementary DLS measurements DLS with a Malvern Zetasizer Ultra by diluting the sample $1 : 10^3$. Since CLiC, NTA, and cryo-TEM measures particle size on the single-particle level, the DLS number average is here used to compare the sizes from the different techniques.

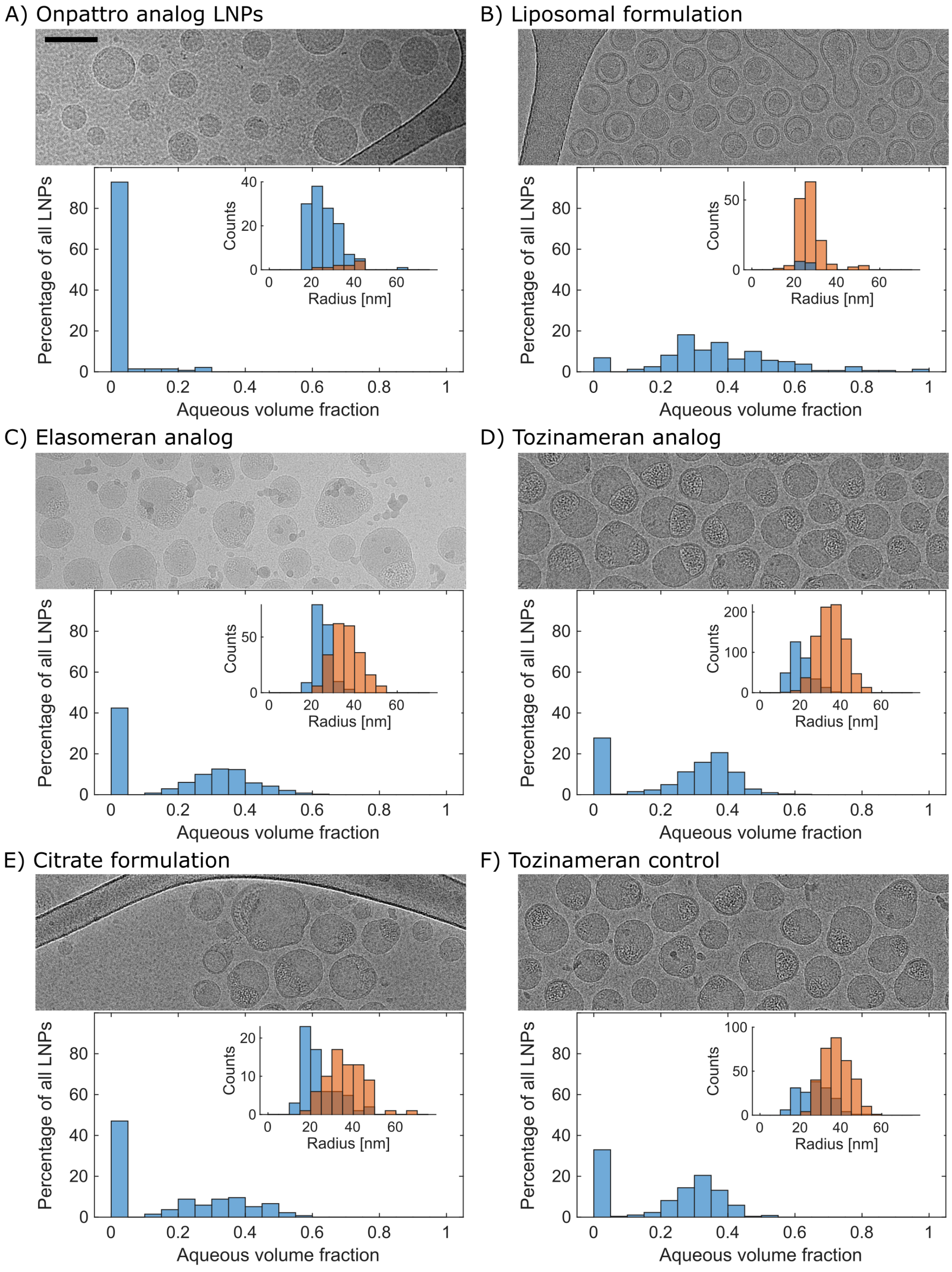

Figure S3: **Structural analysis of cryo-TEM images of LNPs, as a function of formulation parameters.** cryo-TEM images of various formulations were analyzed using the structural classification shown in figure S2. The results are plotted on two histograms for each formulation, including one for the aqueous volume fraction (main) and another for size distributions (insert). The size distributions include subpopulations of LNPs that contain aqueous pockets (orange), and those that don't (blue). Here we observe minimal water content in the Onpattro analog LNPs(A). In contrast, most LNPs in the Liposomal formulation (B) showed significant volume fractions with aqueous content. The rest of the formulations (C - F) showed varying fractions of LNPs with/ or without water content. Scale bar = 100 nm

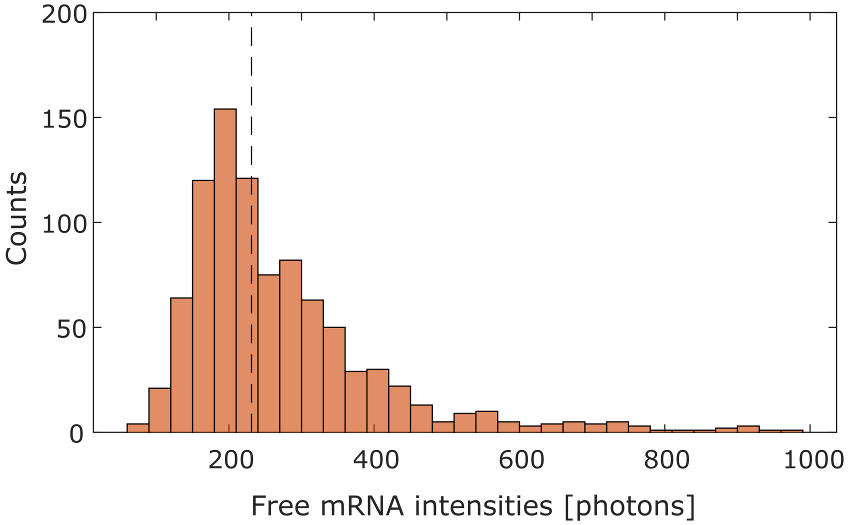

Figure S4: **Distribution of Cy5 labeled free mRNA molecules:** A total of 830 molecules were detected and analyzed. The resulting distribution of single mRNA intensities had a median intensity of 227 photons, with a mean of 263 photons and standard deviation of 130 photons.

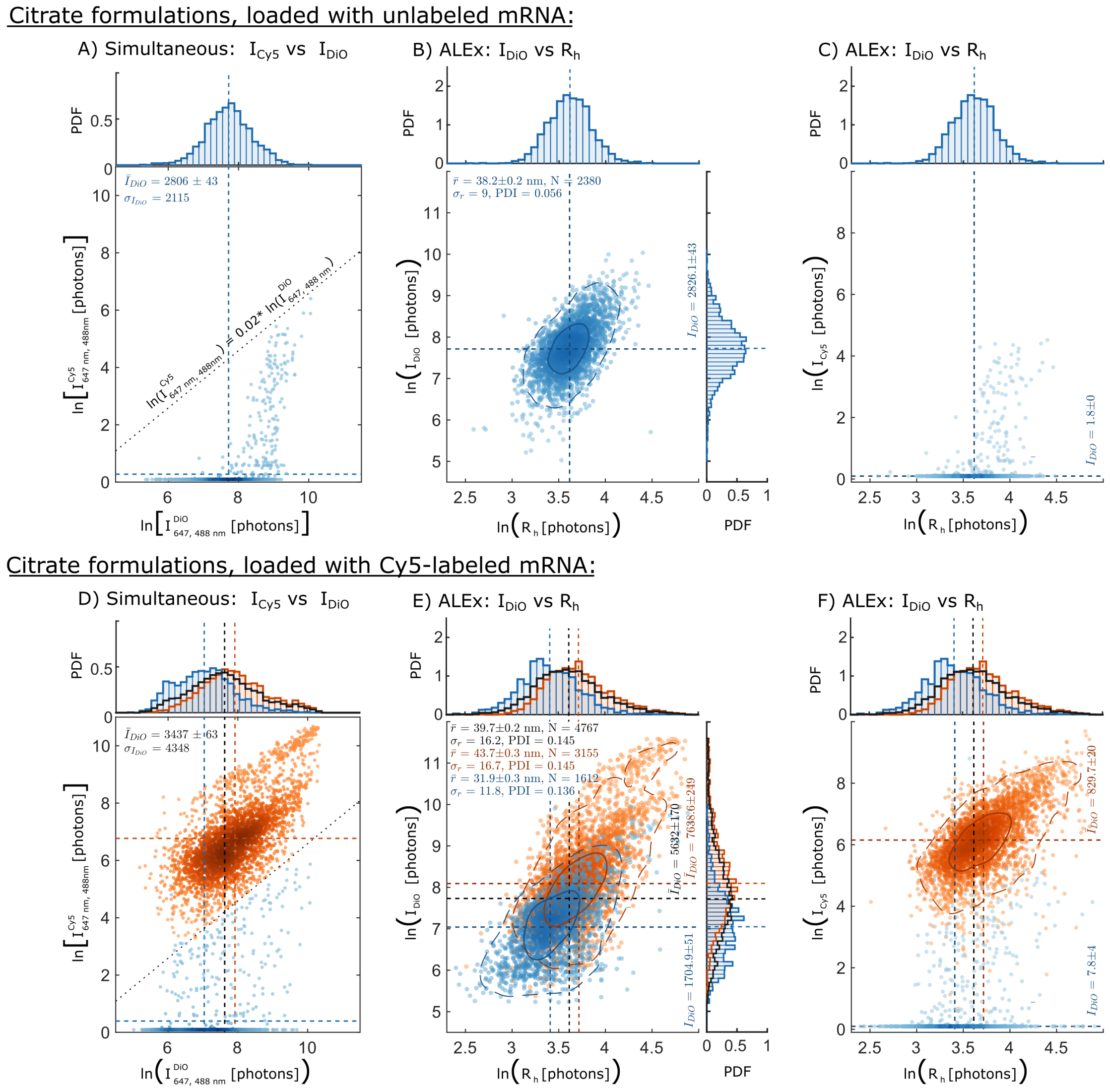

Figure S5: **Measurements on Citrate LNPs with unlabeled mRNA cargo (A - C), versus those with Cy5-mRNA (D - F):** Plots of Cy5 channel intensity ($I^{\mathrm{Cy5}}_{647,488\ \mathrm{nm}}$) versus DiO intensity ($I^{\mathrm{DiO}}_{647,488\ \mathrm{nm}}$), measured with simultaneous excitation settings are shown in (A) and (D). They included a black dashed line that shows the minimum threshold for detecting Cy5 signal, above DiO bleed-through into the Cy5 channel. (B) and (E) show plots of total DiO intensity ($I_{\mathrm{DiO}}$, see equation 1 of main text) versus particle size ($R_{\mathrm{h}}$), for LNPs with unlabeled and labeled mRNA. respectively. Similarly, (C) and (D) show corresponding plots of total Cy5 intensity ($I_{\mathrm{Cy5}}$, see equation 2 of main text) versus particle size ($R_{\mathrm{h}}$). As expected, $\bar{I}^{\mathrm{DiO}}_{647,488\ \mathrm{nm}}$ and $\bar{R}_{\mathrm{h}}$ of control LNPs (vertical lines in A & B) are similar to those of the Citrate formulation (black vertical lines in D and E, respectively). Here black lines represent combined distributions of both empty and loaded LNPs for the Citrate LNPs. Lastly, we observe that the total Cy5 signal ($I_{\mathrm{Cy5}}$) measured in control LNPs (C) have a similar intensity distribution to those of the unloaded subpopulation of Cy5-mRNA LNPs (blue cluster in F). This implies that the signal detected in unloaded LNPs is mostly due to noise levels, and not Cy5 fluorescence.
